# Reconstructing the functional effect of *TP53* somatic mutations on its regulon using causal signalling network modelling

**DOI:** 10.1101/2022.06.23.497293

**Authors:** Charalampos P. Triantafyllidis, Alessandro Barberis, Ana Miar Cuervo, Enio Gjerga, Philip Charlton, Fiona Hartley, Linda Van Bijsterveldt, Julio Saez Rodriguez, Francesca M. Buffa

## Abstract

The gene encoding tumor protein *p53* (*TP53*) is the most frequently mutated gene in human cancer. Mutations in both coding and non-coding regions of *TP53* can disrupt the regulatory function of the transcription factor, but the functional impact of different somatic mutations on the global *TP53* regulon is complex and poorly understood. To address this, we first proceed with a machine learning (ML) approach, and then propose an integrated computational network modelling approach that reconstructs signalling networks using a comprehensive collection of experimental and predicted regulons, and compares their topology. We evaluate both these approaches in a scrutinized pan-cancer analysis of matched genomics and transcriptomics data from 1,457 cell lines (22 cancer types) and 12,531 clinical samples (54 cancer sub-types). Using a ML approach based on penalized generalized linear regression we were able to predict *TP53* mutation, but failed to resolve different mutation types. Thus, to infer the impact of different *TP53* mutations we compared the topological characteristics of the optimized and reconstructed (upwards of twenty thousand) gene networks and extracted gene signatures for each mutation type using network analysis. We demonstrate that by accounting for *TP53* mutation characteristics such as i) mutation type (e.g. missense, nonsense), ii) deleterious consequences of the mutation, or iii) mapping to previously identified hotspots, we can infer a much richer understanding of gene expression regulation, than when simply grouping samples based on their mutation/wild type or gene expression status. Our study highlights a powerful strategy exploiting signalling networks to systematically characterize the functional impact of the full spectrum of somatic mutations. This approach can be applied in general to genetic variation, with clear implications for, but not limited to, the biomedical domain and precision medicine.

## INTRODUCTION

Precise regulation of gene expression is critical to a diverse array of biological processes in health and disease. Dynamic transcriptional changes depending on the physiological circumstances drive cell fate decisions in development, disease, and in response to drugs and mitogens. Transcription factors (TFs) are master regulators of gene expression. Sequence-specific transcription factors can bind to exact regions of the DNA to facilitate transcription initiation of their target genes (Edelman and Fraser [2012]), with this downstream set of genes also known as the *“regulon”*. In cancer, driver mutations have an impact on TFs and many of the key tumor-suppressor genes constitute TFs (Futreal et al. [2004]). Many such mutations lead to structural changes and alter the DNA-binding capacity of the respective transcription factor, impacting their regulatory network/targets. To prevent targeting cancer agents only to a limited set of molecules or mutations and broadening our possibilities for re-purposing of existing drugs, it is critical to understand these networks and to explore the full downstream functional implications of the different mutations.

Gene networks are a powerful tool to study complex diseases such as cancer (Seçilmiş et al. [2020], Reyna et al. [2020], Yan et al. [2016],Zhou [2012], Benstead-Hume et al. [2019], Albert [2005]); however they have not been applied systematically to characterize different mutations. Different variants can be chosen for gene networks, which provide different insights. For example, association networks have been used extensively. While their predictive ability is limited, network metrics (e.g. communities, hubs) can be used to infer gene function (see for example Buffa et al. [2010], Masiero et al. [2013]). Directed networks on the other hand, offer the possibility of representing causality, with inclusion of mode of regulation and differentiation between direct and indirect effects Voukantsis et al. [2019], Melas et al. [2013]. Here, we present a causal network approach to analyse the functional consequence of mutations in TFs, and evaluate its utility to identify the impact that different mutations have on the *TP53* regulatory network (regulon).

*TP53* is central to human biology. The wild-type nuclear tumour protein (p53^WT^), also known as the “Guardian of the Genome”, acts to block cell cycle progression (Agarwal et al. [1995]) in the presence of DNA damage to promote repair or in the case of non-repairable DNA damage, to stimulate programmed cell death through controlling a set of genes required for these processes Levine [2019a]. *TP53* can induce growth arrest or apoptosis depending on the duration and type of stress, the cell type and other physiological circumstances. At present, hundreds of mutations have been identified for the *TP53* gene Levine [2019b] that lead to structural changes that destabilize p53 structure, and alter its DNA-binding capacity and ability to regulate target genes through interaction with transcription regulators and chromatin complexes Kim and Lozano [2018]. It is important to notice that *TP53* mutation status is not always the only determinant, as p53 function can be modulated via alternative mechanisms which impact its upstream regulators. For example, gain or loss of function of MDM2 gene would impact, negatively and positively respectively, on *TP53* stability Chène [2003]. When *TP53* is unable to control the expression of targets involved in tumor repressive responses, it can play a critical role in cancer initiation and progression Suzuki and Matsubara [2011]. While several studies have described p53’s effects on the tumors’ transcriptome and proteome (Donehower et al. [2019], Ozaki and Nakagawara [2011], Muller and Vousden [2014], Mantovani et al. [2019], Lozano [2019], Blagih et al. [2020], Steele and Lane [2005],Klimovich et al. [2019], Joerger and Fersht [2016], Soussi [2000], Melling et al. [2019], Perri et al. [2016]), the implications for the different types of *TP53* mutations are still understudied. Thus, it is timely to take a full advantage of all data available to carry out a systematic analysis of p53 related datasets to come up with ways to predict p53 function in human tumours. Importantly, due to the heterogeneous nature of p53 mutations, distinguish deleterious p53 mutation from other, “passenger”, p53 mutations is of great importance. The differences functional implications between these mutations, could be one of the key reasons why in most cancers TP53 mutation status have not been applied in the clinic to predict patients response to therapy, and to guide clinical practices. Indeed, to be able to link certain types of p53 mutations (missense, deletion or nonsense mutation) with their function is important, and if certain mutation spectrum can be defined as deleterious mutation, it will be very useful.

Using machine learning and network reconstruction methods, and the large number of datasets now available with both whole-transcriptome information and recorded mutation status, we set to study, in a systematic manner, the impact of different *TP53* mutations on the expression of its regulon. To ensure that our reconstructed networks reflect the downstream effect of the real spectra of *TP53* mutations occurring in cancer, and investigate how these impact *TP53* function and clinical outcome, we interrogated 1,457 cell lines across 22 cancer types from the Cancer Cell Line Encyclopedia (CCLE) and 12,531 cancer samples across 54 cancer types and sub-types in The Cancer Genome Atlas (TCGA) databases. First, we assembled a comprehensive list of validated and predicted targets, ranked by level of evidence, using DoRothEA Garcia-Alonso et al. [2019], Garcia-Alonso et al. [2018]. We then followed two approaches: 1) we applied a machine learning approach to build a predictive model of changes in expression of the regulon for different *TP53* mutation types, and 2) we applied a directed gene network approach enabling causal inference to reconstruct *TP53* signalling networks in cancer samples and cell lines harbouring different *TP53* mutations. Analyses of this sort could allow monitoring of the functional impact of specific mutational events in single tumours at diagnosis, or de-novo mutations occurring during treatment, opening new therapeutic options for cancers that are resistant to current therapeutic regimes.

## RESULTS

### Expression of the *TP53* gene is heterogeneous and non-predictive of mutation status

The *TP53* mutational landscape across human cancers and cell lines is very heterogeneous (see Willis et al. [2004], Olivier et al. [2010], Olivier et al. [2010], Petitjean et al. [2007], Shahbandi et al. [2020]. In our analysis we found that *TP53* was mutated in 4,250 / 12,531 (34%) of the TCGA cancer samples and 898 / 1,457 (61%) of cell lines, with some cancer types showing strong mutational frequency (such as ovarian, lung and glioblastoma) and others much less (Suppl. Figure 1(a), 1(b). Approximately 65% of all found mutations were missense mutations (point mutations where a single nucleotide change codes for a different amino acid) in both TCGA and CCLE (Suppl. Figure 1(c), 1(d)). This is in agreement with previous reports (Donehower et al. [2019], Kotler et al. [2018], Tan and Luo [2009], Walerych et al. [2016]).

Mutation type and deleterious function of mutation (a deleterious mutation is a mutation that renders the protein non-functional) respectively were extracted from the CCLE and TCGA annotation, and we considered known hotspot mutations with protein changes as follows: p.R175H, p.R248Q, p.R273H, p.R248W, p.R273C, p.R282W, p.G245S Baugh et al. [2017]. These were all characterized as missense mutations. Finally, both analysis of TCGA tumor samples and CCLE cell line data show that only 10% of samples contain more than a single mutation for *TP53*.

We then asked how these different mutations correlated with gene expression of TP53. Expression varied across different mutations, with some mutations resulting in higher expression with respect to WT status, and others lower. Interestingly we observed a very high concordance between cell lines and cancer samples. Samples with frame-shift mutations (deletions/insertions) and nonsense mutations showed generally lower expression than WT samples in both CCLE and TCGA. In-frame (deletions/insertions) and missense mutations showed higher levels of expression, while other mutations showed levels of expression similar to WT (Suppl. Figures 2(a),2(b)). Samples with deleterious mutations and hotspot mutations showed respectively a significantly lower (Suppl. Figures 2(d),2(f)) and higher (Suppl. Figures 2(c), 2(e)) expression of *TP53* than other mutations.

We also looked at Copy Number Variation (CNV) (Suppl. Figure 3). In CCLE, copy number gain was associated with higher median expression across mutated samples, whereas diploidy projects higher expression in wild-type cell lines. In TCGA, the highest expression in mutated tumour samples was found in amplification, whereas besides deep and shallow deletions in wild-type samples, all other types of copy number variation had similar but higher expression values.

While carcinogenesis implies bi-allelic loss of functional tumor suppressor genes, the most typical *TP53* mutation configuration is a single *TP53* mutation with loss of the remaining *TP53* allele through a large-scale deletion on chromosome 17p (Nigro et al. [1989], Baker et al. [1990]). Additionally, mutant p53 can have a dominant negative effect over *p53^WT^* and/or gain of function activity independently of the wild-type protein Willis et al. [2004]. Furthermore, data from the TCGA and CCLE (Donehower et al. [2019]), show that only 10% of samples contain more than a single mutation for TP53. There is also evidence that single mutations of *TP53* are associated with loss of a single allele (2/3 of tumors) and a high distortion CNV, whereas tumors with more than two mutations usually retain both alleles (diploid in almost one third of cases), Donehower et al. [2019].

### Association with prognosis supports functional heterogeneity of *TP53* mutations

*TP53* mutations have been associated with poor prognosis Kandoth et al. [2013]; however, the landscape is complex (for a review see e.g. Robles and Harris [2010]). In some cancers, such as hematological malignancies or ovarian cancer, *TP53* status is also used to guide the treatment strategy. In others, such as breast cancer, some evidence of association of *TP53* mutation with poor survival have been produced, however the landscape is not entirely clear Shahbandi et al. [2020], and it is largely dependent on treatment and accurate characterization of variant classification (mutation type). Some specific mutations can have a significantly different impact on survival than others; for example, missense mutations within exons 5 to 8, corresponding to the DNA binding domain, have been show to correlate with poorer survival than silent mutations in this region or no mutations Olivier et al. [2009].

Building on this evidence, we asked whether stratifying patients by any of the different types of p53 mutation would allow us to better correlate *TP53* with patient disease outcome than simply grouping patients by wild-type or mutant status. We interrogated TCGA data-sets and considered the different sample stratification: i) samples with *TP53* mutant versus WT status, ii) samples carrying missense mutations vs other mutations (excluding WT samples), iii) samples carrying missense mutations vs other mutations vs WT (3 groups), iv) samples with deleterious mutations vs non-deleterious mutations vs WT (3 groups), v) samples carrying hotspot mutations vs other mutations vs WT (3 groups) (Suppl. Figure 4).

The results show that *TP53* mutation was associated with worse prognosis (Suppl. Figure 4(a)). Furthermore, samples carrying missense mutations had worse prognosis than samples with non-missense mutations (Suppl. Figure4(b)), even when mixing with WT samples (Suppl. Figure 4(c)), and samples with hot-spot mutations had worse prognosis than samples with non-hotspot mutations and WT groups (Suppl. Figure 4(e)). Deleterious mutations also showed worse prognosis to non-deleterious mutations and WT groups (Suppl. Figure 4(d)). Furthermore, specifically for missense mutations when comparing the deleterious versus the non-deleterious we also see worse prognosis for the non-deleterious (Suppl. Figure 4(f)). This is in agreement with previous research Donehower et al. [2019] as a missense mutation, changes the structure of the p53 protein but also makes the protein negative dominant on the WT version (which is a tumor suppressor). Finally, when comparing hotspot and non-hotspot but both deleterious, the worse prognosis is seen in the case of non-hotspot deleterious cases (Suppl. Figure 4(g)). These results taken together confirm that *TP53* mutational status correlates with clinical outcome and importantly, that specific types of mutations affect patient survival differently. Furthermore, we compared the survival curves in two additional settings in relation to TP53 hotspot mutations (as those appear in Baugh et al. [2017]) in Supp. Figure 5. We can see in Suppl. Figure 5(a) that when we include the WT samples, we clearly see a difference in survival (p « 0.05) but as shown in Supp. Figure 5(b), by removing the WT samples from the data this difference is no longer significant.

### A ML classifier based on expression of *TP53* regulon predicts *TP53* mutation status (WT/MT), but not the type of the mutation

An initial principal component analysis (PCA) of the mRNA expression of the regulatory network of *TP53* (regulon) indicated a significant variation across samples (Suppl. Figure 6, principal component one accounting for 15.6% of the variation while the second equivalently for 11.5%), which suggests differential regulation of these genes, but not a global association with mutation. A basic clustering analysis confirmed these results (Suppl. Figure 7) showing different patterns of up or down-regulation for different groups of regulon genes, but not an immediately clear correlation with mutation status. Furthermore, a previous basic signature of four genes (CDC20,CENPA,KIF2C,PLK1) showed a significant positive correlation with the presence of p53 mutation in clinical samples (Donehower et al. [2019]), in other words these four genes had differential expression levels in *TP53* mutated samples with respect to normal.

These results together indicate that although the expression of *TP53* itself is not predictive of mutation status, specific gene expression features in the clinical samples could be. However, it is not easy to draw conclusions from this study as these genes were not validated targets of *TP53* nor were they developed into a predictor. On the other hand, these four genes were characterized by higher expression in TP53 mutated samples with respect to WT TP53, suggesting that TP53 represses their expression when active. Many of the repressed genes do not contain binding sites hence they tend to be not represented in regulon databases, and this could explain why they have not been validated as target genes in previous studies. So, we asked first if we could develop a predictor for these 4 genes, and then how this performed with respect to a predictor based on *TP53* deposited targets. Finally, we asked if these gene signatures could predict not only mutation status but also the type of mutation present in the specific sample/group of samples.

To develop a predictor of *TP53* mutational status based on gene expression we used a machine learning (ML) approach based on generalized linear models and Penalized Regression (see Methods). This method has performed well in our hands before to reconstruct gene networks, and given the TF network will have a large number of directly regulated targets, we can hypothesize that a linear regression captures this signal sufficiently well. First, we considered the four genes previously identified and the *TP53* regulon, with the following aims: i) prediction of missense mutation with respect to other mutations, ii) prediction of any *TP53* mutation, and iii) prediction of hotspot mutations. We did this in a cross-validation setting. To achieve this, we applied cross-validation and varied the train-test set proportions during the re-sampling to ensure that the size of the training dataset did not affect significantly the results (see Methods).

We observed that both the previously identified set of 4 genes and a comprehensive regulon signature could predict well the presence of *TP53* mutation of any type, in both the cell line data (Figure 1(d),1(e)) and cancer samples (Figure 2(c),2(d)). This confirms, as expected, that the regulon usage is indeed generally different in *TP53* mutant tumours with respect to *TP53* wilde type. Specifically, a 4 gene model showed an average optimal misclassification of 15% and a model built with the extended set of regulon genes showed an average of 0% miss-classification at the optimal condition, suggesting that the use of the full regulon provides an advantage with respect to using only 4 genes. When we tried to predict the type of mutations (missense vs non-missense/any mutation) the models’ performance deteriorated (Figures 1(b),1(c), 2(a),2(b)). The average misclassification rate was between 35 and 40% for CCLE data and above 40% for TCGA data. Eventually, using this approach we could not build a predictive classifier of the type of *TP53* mutations. Thus, we could not detect significant global consequences on the expression measured as RNA levels of the *TP53* regulon, in samples with different types of *TP53* mutations. Importantly, these differences might be there, but not strong or stable enough to allow a robust predictor to be built. Additionally, we attempted to calculate a binomial classifier between the hotspot mutations of TP53 (as those appear in Baugh et al. [2017]) and any other mutation in both CCLE and TCGA. In this case the signal was very good and we could detected differences in the regulon usage. Both of the signatures perform well in CCLE cell lines 1(f),1(g), averaging around 10% miss-classification errors, whilst in the TCGA case the 4-gene signature performs better than the regulon (less than 20% miss-classification error as opposed to approximately 30% for the regulon) Figure 2(e),2(f).

**Figure 1:**
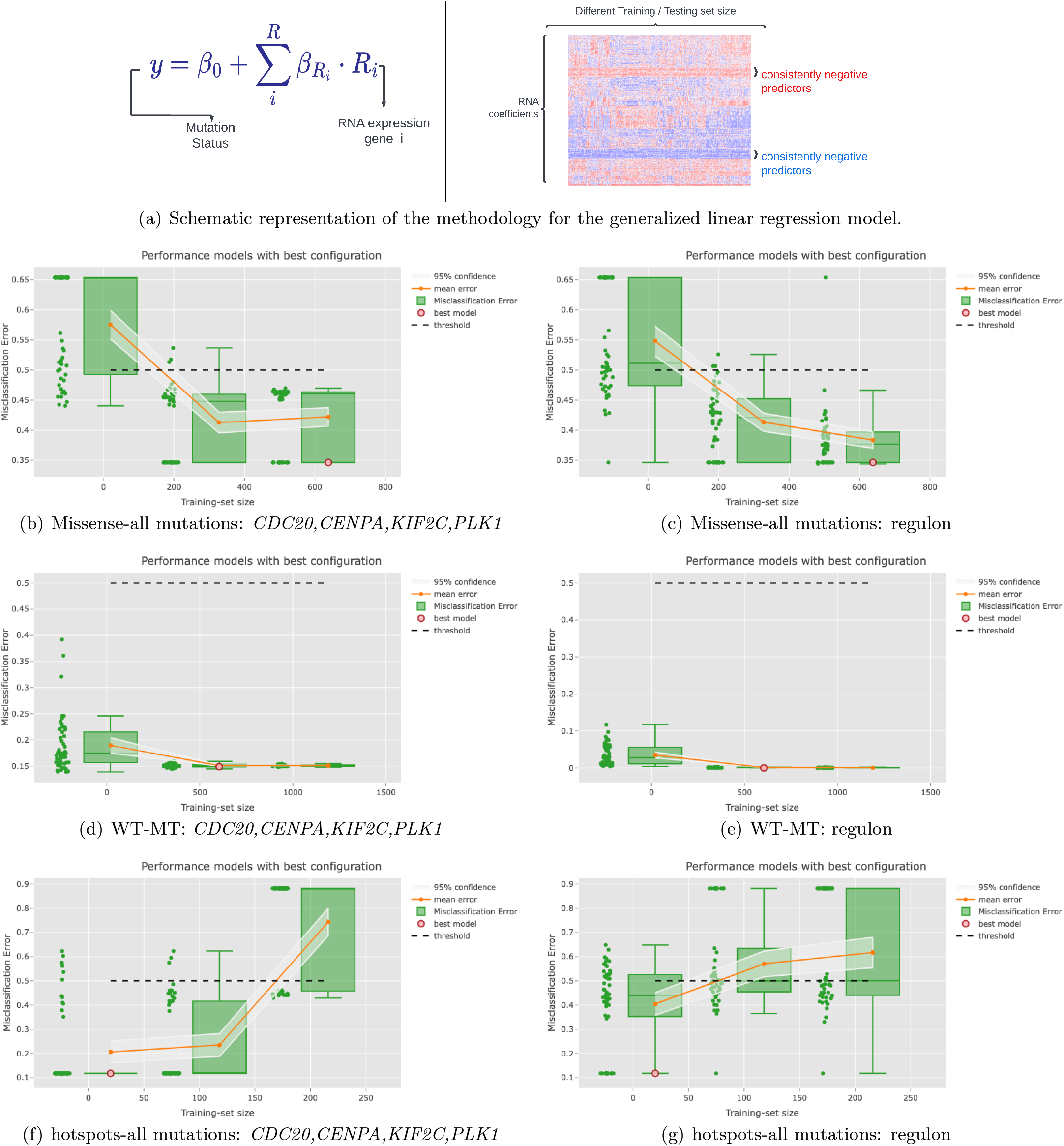
Comparing a model built using a comprehensive regulon with a model developed using a minimal 4-gene signature in the CCLE dataset. Penalized regression was used with multiple settings. The models were trained to predict three different p53 mutational features: (a) Briefly an elastic-net model (see methods) is built in cross-validation for different train-test combinations and misclassification error assessed. We can also extract and compare the gene frequently occurring in the models, missense mutations (b-c), or simply any mutations (d-e) and finally hotspot p53 mutations (f-g). No WT samples were included besides in (d-e). In each plot, the x-axis represents the different training set sizes while the y-axis shows the accuracy measure (i.e. the misclassification error) used to assess the performance of the fitted models. The mean error and the associated confidence interval are also reported for each training set size. Each green dot in the plots corresponds to a trained model. The red dot represents the best model selected in cross-validation (see Methods). Different training set sizes are used, and the one providing prediction error with the lowest upper confidence interval was chosen. The best model is then selected so as to have the minimum misclassification error. The figures indicate the misclassification errors in the full-data set.

**Figure 2:**
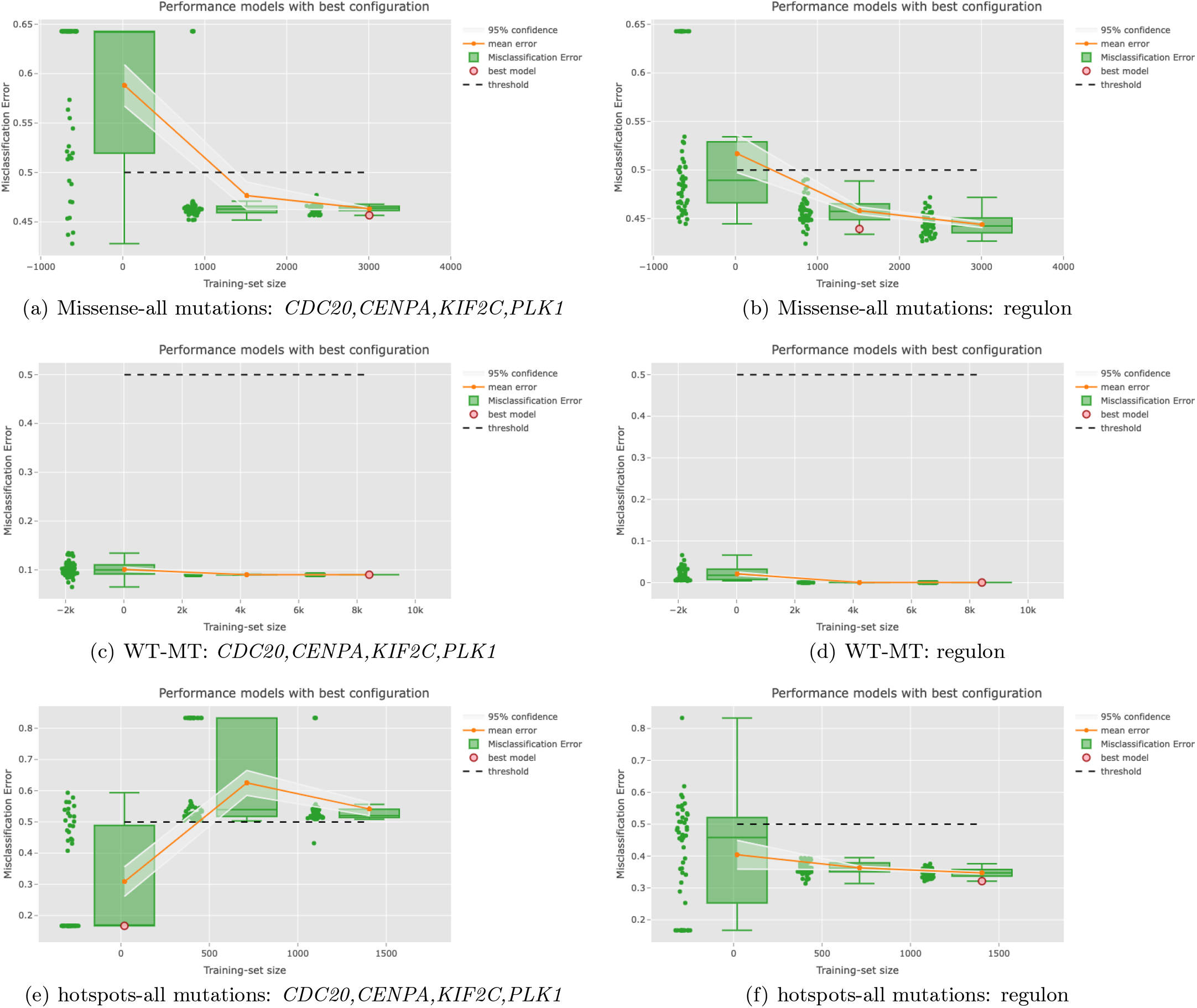
Similar as Figure 1 but for TCGA. Again we compare across (a-b) Missense versus all other types of mutations (no WT samples included), (c-d) WT versus any mutation and (e-f) hotspot p53 mutations versus all other non-hotspot mutations (again no WT samples included). We can see that the 4-gene signature behaves worse than the regulon signature across all settings in terms of misclassification error in the full data set.

### *TP53* mutations of the same type, with deleterious function and similar protein changes show network similarity in cell lines and cancer samples

To investigate the different functional implications of different types of *TP53* mutations at a global scale, we asked how the different mutations correlated with expression not only of the targets (the regulon) of TP53, but also the expression of genes in the upstream and downstream, but indirect, network. So, we asked if we could observe global network rewiring in TP53 mutant cases, and in different mutation types. To address this we used directed edges network modelling. First, we assembled a Prior Knowledge Network (PKN) for *TP53* regulon using OmniPath Turei et al. [2016] for the human genome. We considered only targets with the known mode of regulation and where the source to target information was available. We then processed the expression matrices from CCLE and TCGA to include expression of all these target genes. The resulting PKN is a signalling network including approximately 50k signed interactions (1: up-regulation, −1: down-regulation). Since we are interested in the downstream effect of *TP53* transcription factor, we used the CARNIVAL pipeline to prune edges from this large pool of interactions. We kept only those edges which emanate from TP53, directly or indirectly, meaning there exists a path interconnecting *TP53* with the gene each time.

Next, we defined the perturbation of *TP53* based on its deleterious status as a knock down when true, and as active when false (based on CCLE and TCGA Deleterious feature annotation for each single mutation). For the causal gene network modelling inference we used CARNIVAL in combination with the CPLEX optimization studio. CARNIVAL requires a triplet to be initialized: i) Sign of perturbation (activated or not), ii) Prior Knowledge Network and iii) Expression matrix of the regulon of the TF.

First we define the transcription factor of interest (TP53 in this case) and its status as derived by the deleterious function of the specified mutation in each single sample (CARNIVAL works on single samples). The expression matrix is converted to Normalized Enrichment Scores (NES) per sample (using the VIPER package). We then reconstruct the topology and gene activity profile for the regulon by minimizing the mismatch between the predicted state of each gene (according to the consistency rules imposed by the optimization model’s constraints) and the NES scores. This reconstruction changes effectively the topology of the network as well as the genes that are involved (removed or added). For a more detailed description of the Mixed-Integer Linear Optimization Model (MILP) which underpins CARNIVAL machinery see Methods.

Once each network has been optimized and thus reconstructed, we then compared the networks calculating a network similarity score between all pairs of optimized networks generated for each *TP53* mutation appearing on any sample in the CCLE and TCGA mutations profile matrices. For details on how we derive the network similarity see Methods. It is important to clarify that in Table 1 the first four columns include the number of all possible combinations of pairs of networks and thus the total number of comparisons induced is maximized, whereas in the rest columns we move to a *conditional universe* since we filter for specific *TP53* statuses; thus the number of total comparisons shrinks compared to the prior (general) pool. However, the percentage comparisons eventually phrase the results proportionally to the respective pool of total comparisons in each studied case.

**Table 1:** This table summarizes the CARNIVAL results for all 22 different types of cancer tested in CCLE. We report on the similarity of the optimized networks across four different settings (general, mutation type,deleterious function, hotspots) and similarity percentages (25,50,75 and 90%).

|  | CCLE Cancer | G25 | G50 | G75 | G90 | M25 | M50 | M75 | M90 | D25 | D50 | D75 | D90 | H25 | H50 | H75 | H90 |
| --- | --- | --- | --- | --- | --- | --- | --- | --- | --- | --- | --- | --- | --- | --- | --- | --- | --- |
| 1 | bone | 83 | 52 | 24 | 3 | 100 | 97 | 41 | 8 | 100 | 96 | 44 | 6 | 100 | 100 | 17 | 0 |
| 2 | breast | 61 | 45 | 9 | 0 | 100 | 86 | 19 | 1 | 100 | 88 | 18 | 1 | 100 | 62 | 19 | 0 |
| 3 | colorectal | 79 | 53 | 28 | 2 | 100 | 98 | 47 | 4 | 99 | 96 | 49 | 4 | 100 | 95 | 32 | 3 |
| 4 | esophagus | 68 | 49 | 13 | 1 | 100 | 92 | 21 | 2 | 100 | 93 | 24 | 2 | 100 | 100 | 0 | 0 |
| 5 | gastric | 75 | 51 | 9 | 1 | 98 | 79 | 14 | 1 | 98 | 80 | 14 | 1 | 100 | 100 | 30 | 0 |
| 6 | glioblastoma | 94 | 85 | 41 | 2 | 100 | 98 | 43 | 3 | 100 | 98 | 48 | 2 | NA | NA | NA | NA |
| 7 | glioma | 85 | 72 | 34 | 2 | 100 | 99 | 45 | 4 | 100 | 100 | 47 | 3 | 100 | 100 | 38 | 5 |
| 8 | kidney | 63 | 45 | 12 | 3 | 100 | 92 | 12 | 8 | 100 | 96 | 27 | 6 | NA | NA | NA | NA |
| 9 | leukemia | 70 | 52 | 33 | 5 | 98 | 96 | 66 | 11 | 98 | 97 | 63 | 10 | 100 | 100 | 100 | 7 |
| 10 | liver | 60 | 49 | 12 | 1 | 100 | 92 | 25 | 2 | 100 | 92 | 23 | 2 | NA | NA | NA | NA |
| 11 | lung | 62 | 43 | 8 | 0 | 100 | 76 | 14 | 1 | 100 | 78 | 15 | 1 | 100 | 67 | 11 | 0 |
| 12 | lung NSC | 65 | 48 | 12 | 1 | 100 | 84 | 20 | 1 | 100 | 86 | 21 | 1 | 100 | 57 | 10 | 0 |
| 13 | lung small | 66 | 44 | 11 | 0 | 100 | 83 | 21 | 1 | 100 | 82 | 20 | 1 | 100 | 100 | 0 | 0 |
| 14 | medulloblastoma | 73 | 40 | 13 | 7 | 73 | 40 | 13 | 7 | 73 | 40 | 13 | 7 | NA | NA | NA | NA |
| 15 | melanoma | 64 | 46 | 15 | 3 | 100 | 85 | 29 | 5 | 100 | 87 | 28 | 5 | NA | NA | NA | NA |
| 16 | ovary | 61 | 42 | 6 | 0 | 100 | 76 | 8 | 1 | 99 | 78 | 11 | 1 | 100 | 80 | 7 | 0 |
| 17 | pancreas | 68 | 49 | 11 | 1 | 100 | 85 | 16 | 1 | 100 | 87 | 19 | 1 | 100 | 100 | 0 | 0 |
| 18 | prostate | 50 | 50 | 17 | 17 | 100 | 100 | 0 | 0 | 100 | 100 | 33 | 33 | NA | NA | NA | NA |
| 19 | skin | 67 | 48 | 12 | 2 | 100 | 87 | 20 | 4 | 100 | 89 | 22 | 4 | 100 | 100 | 100 | 0 |
| 20 | thyroid | 82 | 63 | 13 | 1 | 100 | 95 | 20 | 2 | 100 | 96 | 20 | 1 | 100 | 100 | 33 | 0 |
| 21 | TNBC | 62 | 45 | 8 | 1 | 100 | 84 | 17 | 0 | 99 | 86 | 16 | 1 | 100 | 50 | 17 | 0 |
| 22 | urinary tract | 66 | 54 | 10 | 2 | 97 | 85 | 13 | 2 | 95 | 84 | 15 | 2 | 100 | 67 | 0 | 0 |

**Table 2:**
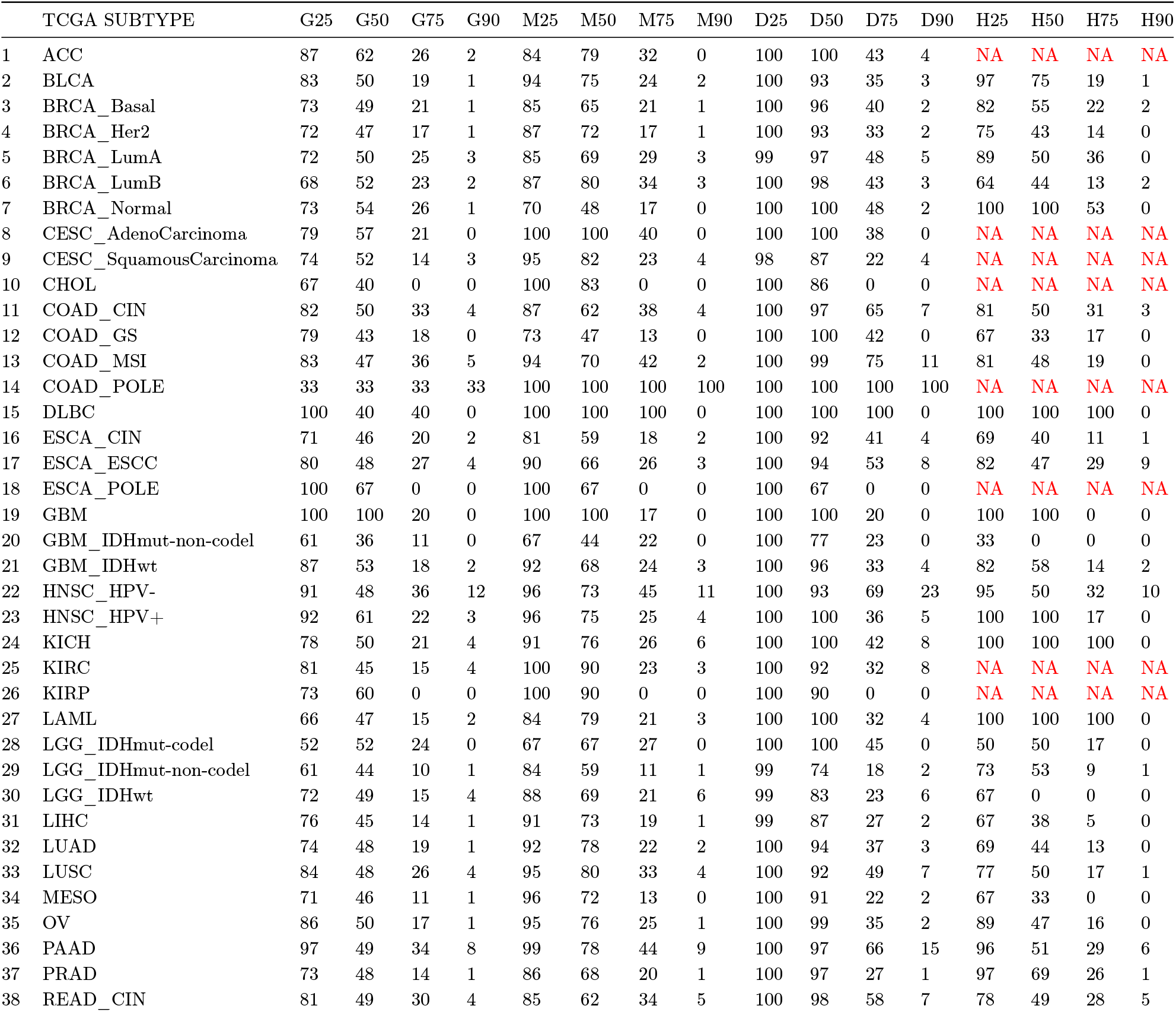
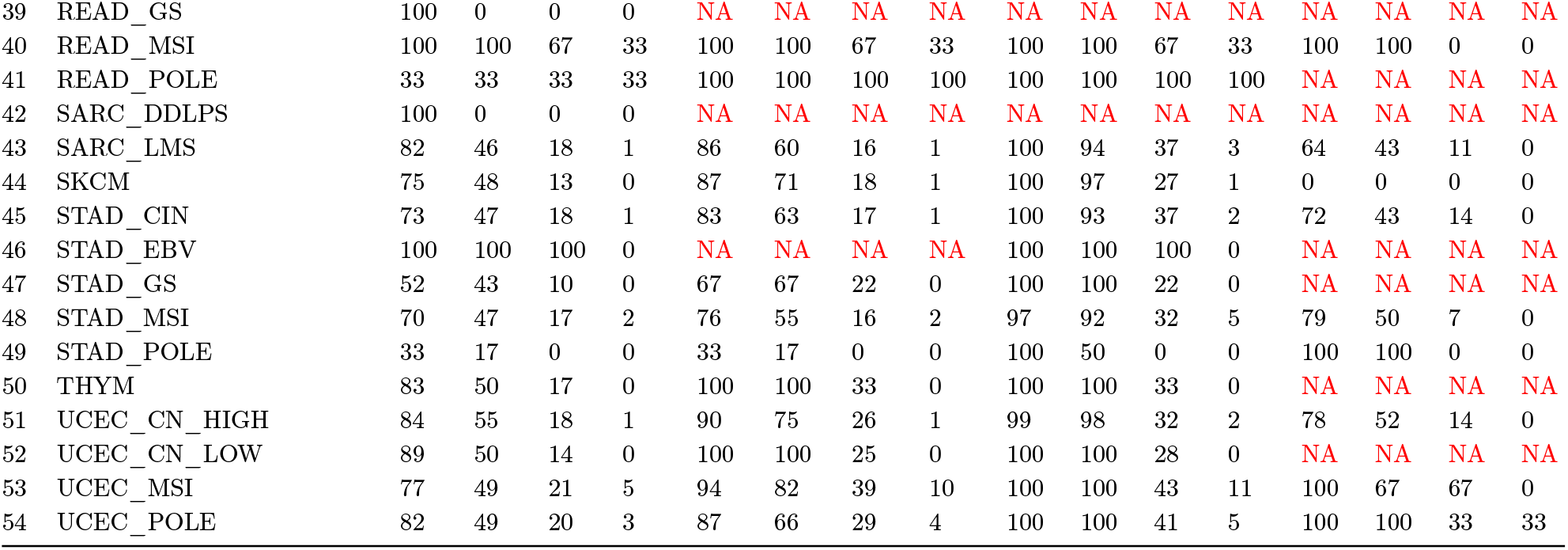
The CARNIVAL results on 54 different sub-types of cancer in TCGA, across different scores of similarity and different metrics such as mutation type or deleterious function of the gene.

Supplementary Table 1 contains the full results per cancer type, or pan-cancer across all four different settings (columns) and four percentages of similarity measured (25, 50, 75 and 90%) for CCLE 21Q2 and Suppl. Table 2 similarly for 54 sub-types in TCGA. Figures 4,5 visualize the respective tables using radar plots. In Suppl. Figure 8 we see three plots for entirely different comparison scenarios: in (a) we see projected similarity when we compare networks that only have different mutation type, (b) is the same but for the deleterious function impact of the mutation on the TF and in (c) we compare networks that the TF mutation corresponds to an identical protein change. These tests taken together show the consistency of the inference, that is, the criterion for which we compared the networks projects high similarity not at random but at a functional relationship level.

**Figure 3:**
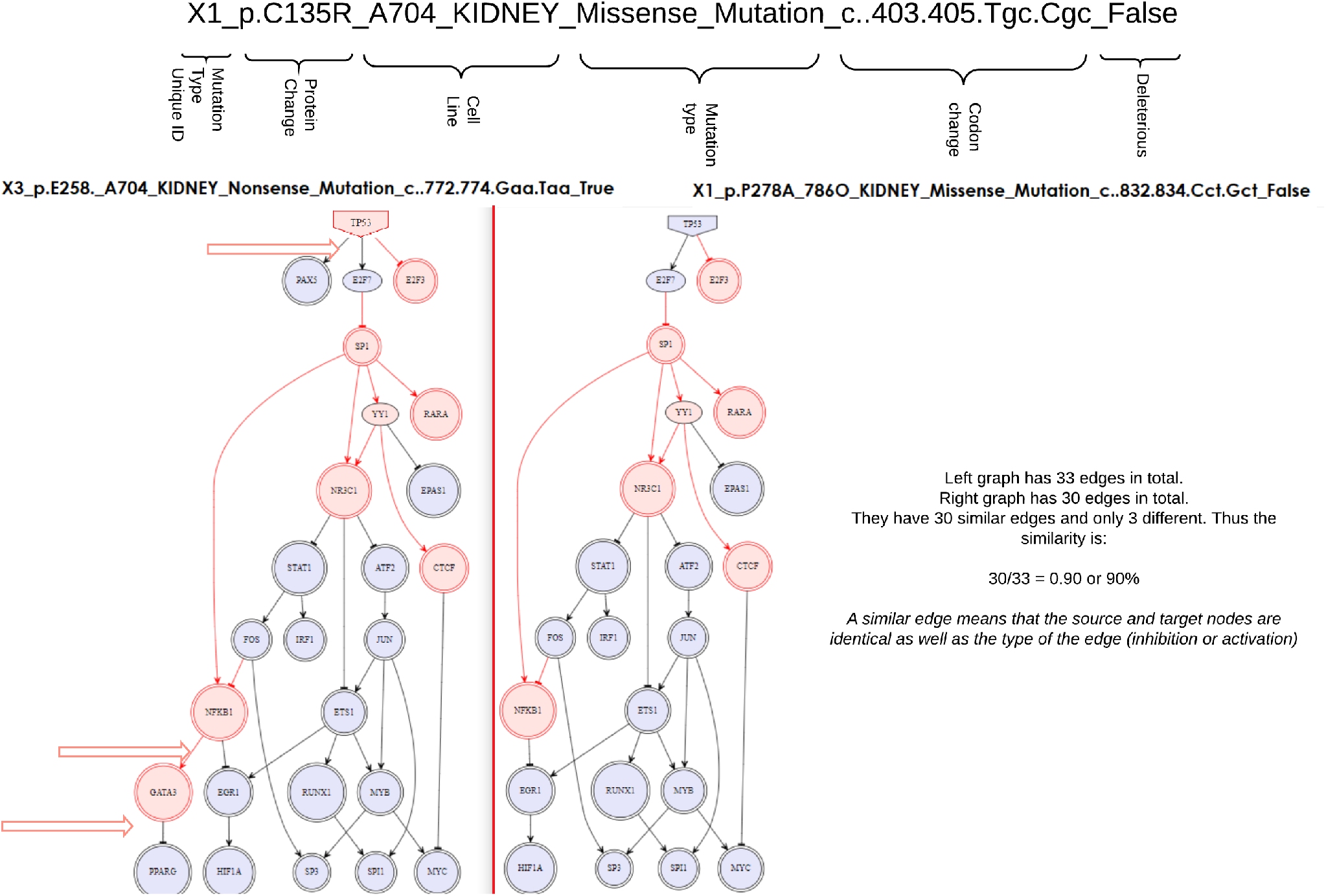
Illustration of our network comparison approach is exemplified using reconstructed networks in two distinct samples from kidney cancer. A different perturbation results to a different downstream effect which is captured by the percentage of similarity between the two networks/graphs.

**Figure 4:**
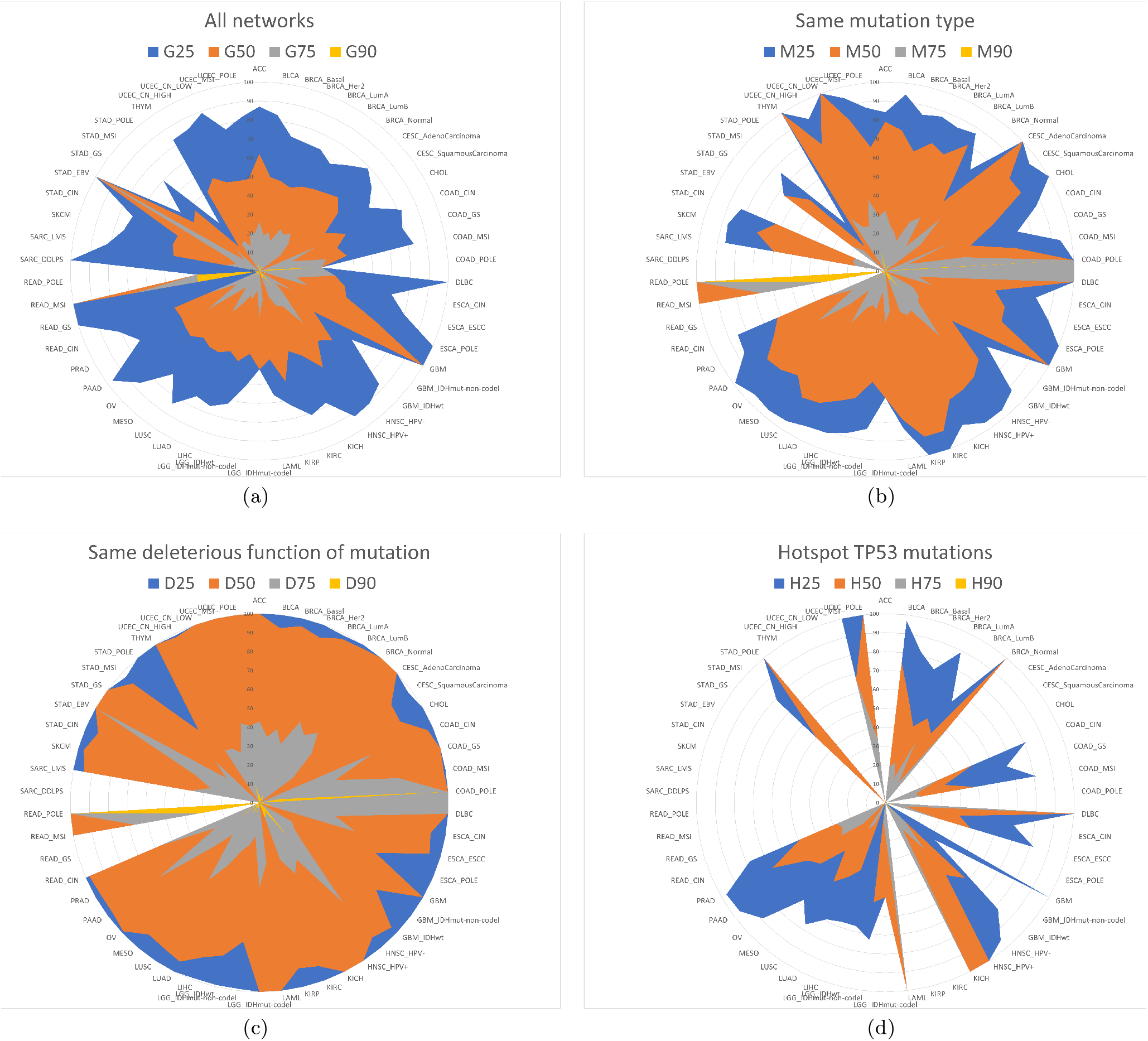
TCGA per sub-type results for all p53 mutations optimized using CARNIVAL. We see that the similarity score (blue, red, grey, yellow correspondingly for 25, 50, 75 an 90%) is higher (the lines expand towards the outer perimeter) when we filter the comparison of the optimized signalling networks by i) mutation type (b) and ii) deleterious function of the mutation for the gene (c), as opposed to the general case in (a).

**Figure 5:**
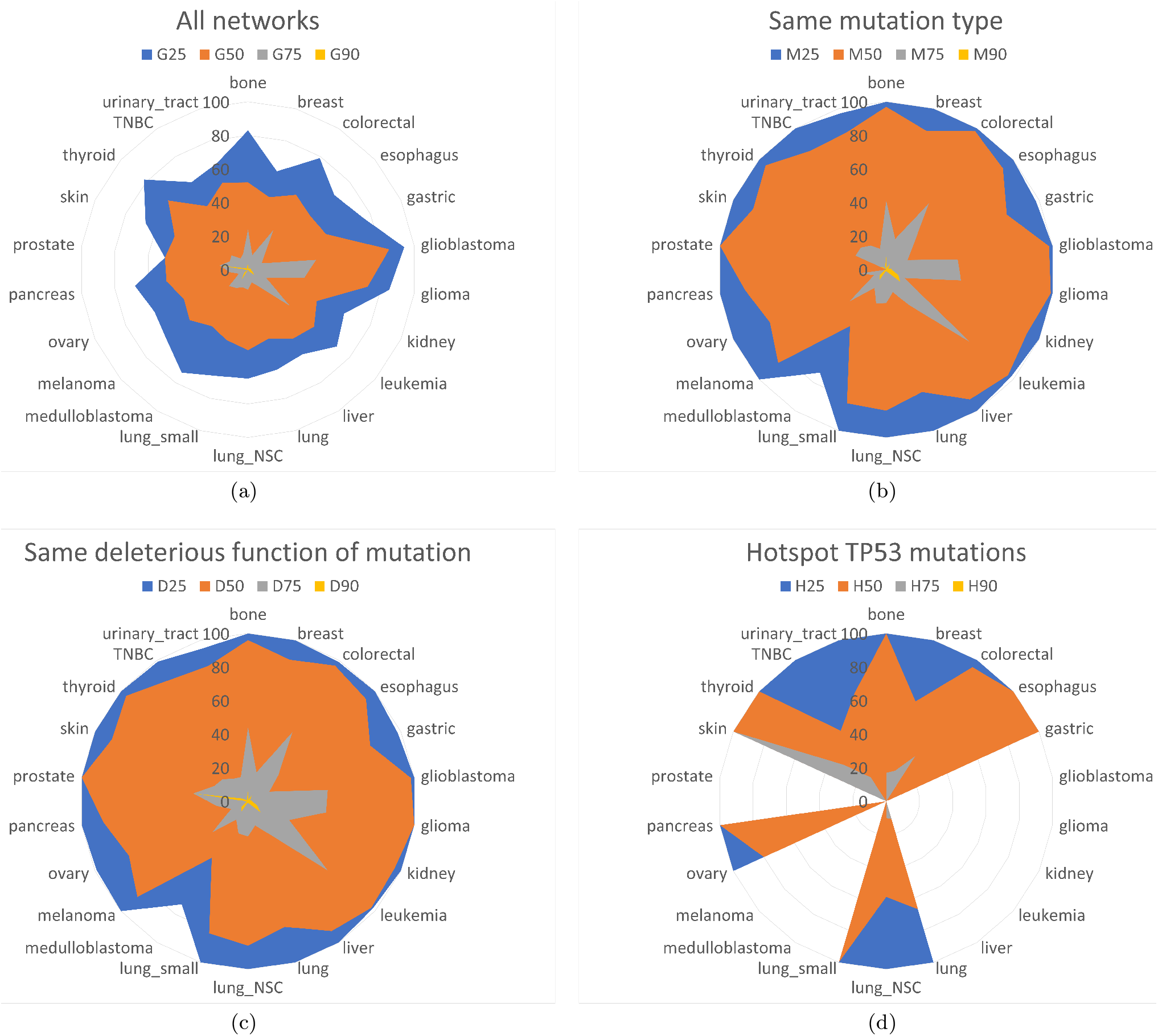
The above four radar/spider plots summarize the main results emanating from the full computational study performed across CCLE. The radar/spider plots indicate the percentages of similarity in the corresponding optimized *TP53* mutation induced downstream regulon across four different metrics (each metric corresponds to one of the four given graphs). Across multiple cancer types labeled on the perimeter of the outer circle (cell lines in CCLE), we compare the CARNIVAL optimized networks on the basis of four different metrics (colors) as shown in the figure. The networks have been optimized such as they fit the expression data (internally translated to Normalised Enrichment Scores - NES via VIPER package) input each time for the Prior Knowledge Network (PKN). We then rank their similarity score in each radar plot (homo-centric circles ranging in [0,100]; see Methods for an explanation of the quantification of similarity in this context). We can see that once we filter the mutations per type (ii) or deleterious function (iii) for *TP53*, networks visibly become more similar in contrast with the general setting in case (i). Hotspot mutation similarity also remains strong, but the lack of occurrence in some cancers might be skewing the overall picture.

The first two tests re-affirm that the mutation type and the deleterious impact of the mutation for the gene play a strongly important role for the downstream signal in the way the TF interacts with its regulon. The last test shows that when the pair of compared networks come from a mutation of the TF which corresponds to the same protein change we also get very similar network topology.

According to our analyses, the similarity percentage in both CCLE and TCGA significantly improves across all levels compared (25,50,75 and 90%), when we move from general/total similarity (all possible combinations of networks compared without filtering for a specific p53 feature) to either same type of mutation or deleterious functional impact of the mutation for TP53. It is evident from the radar plots in Figures 4,5 that the blue (general 25%) similarity curve expands almost to the perimeter of the plot (thus reaching higher score/percentage) when we move to the same type of mutation as opposed to the total similarity on the first radar plot. This continues for the orange (50%) similarity curve and gradually decrease as we become more stringent on the similarity percentage sought (yellow and grey curves, correspondingly for 75 and 90% and above, similarity). The same effect is observed in the case of deleterious filtering whereas for the hotspots case the results show a lesser signal of association between the feature and the downstream graph similarity on the optimized network, which though might be an effect of fewer observed cell lines and tumour samples harbouring one of the analyzed hotspot mutations (as per the hotspots identified in Baugh et al. [2017]).

More specifically, in the case of TCGA, 36/54 sub-types have a greater percentage of networks reaching the 75% cutoff similarity threshold when only cases with the same type of mutation are included than in the general case. This ratio improves when we compare against same deleterious feature of mutation to 45/54.

In the same way, in CCLE21Q2 19/22 cancer types improve the total number of similar networks when we set the cutoff similarity threshold to 75% and above and comparing the general case with same type of mutation. This also improves when we compare against the same deleterious feature of mutation in 21/22 cancer types. For CCLE, Leukemia sees the biggest similarity gain moving from the general case to the same type of mutation (up 33%).

### TP53 perturbation using radiation and hypoxic stress is associated with differential usage of its regulon

#### Network modelling on WT *TP53* lung cancer cell lines under radiation reveals different downstream effects on its regulon based on p53 perturbation as an after effect of irradiation

We investigated the downstream effect of different perturbations of p53 using WT *TP53* cell lines in lung cancer (H460) between no treatment (0h) and after treating with radiation. The experiment is a time course (0 = pre-radiation, and 2, 6, 12, 24hrs post 2Gy radiation) with 3 cell lines (H460 Parental, and 2 radio-resistant - *H*460_50B and *H*460_60A). Suppl. Figures 9,10 shows the computational results after the optimization of the respective different conditions. The results indicate that after irradiation (>0h) p53 is activated and thus the signal downstream changes drastically in comparison to samples before irradiation (where p53 is inactive). In Supp. Figure 10 we can clearly see that once we compare only irradiated with non-irradiated cell-lines, the network similarity diminishes to 0% for similarity greater than 50% across the two compared reconstructed (optimized) graphs/networks.

#### Breast cancer cell line experiments under normoxia and hypoxia reveal the downstream regulating effect of *TP53* is vastly different based on condition

We also investigated the effect of hypoxia on *TP53* using RNA-sequencing data from two independent experiments. In the first experiment, a panel of four breast cancer cell lines (MCF7; MDA-MB-231; HCC1806; MDA-MB-468) were exposed to 1% or 0.1% *O_2_* and collected at 24hr and 48hr time-points. The second, smaller experiment (45 samples) focused on only two of these cell lines (MCF7 and MDA-MB-231), and was limited to a single hypoxia condition (24hr, 1% O_2_). In the second and larger (360 samples) experiment we found that the percentage of networks with similarity greater or equal than 25, 50, 75, 90% was 100%, 55%, 53% and 30%; when we filtered for inverse conditions (hypoxia/normoxia) we saw the similarity reduced dramatically to 100%, 0%, 0%, 0%.

When we compared the 45 networks from the smaller experiment we found that the percentage of networks with similarity greater or equal than 25, 50, 75, 90% was found to be 100%, 49%, 36% and 25% respectively (in the general random comparison pooling case). Filtering the comparisons only between samples that correspond to inverse oxygen conditions, that is either 0.1 or 1% hypoxia on one side and normoxia on the other, we found that the percentage of networks with similarity greater or equal than 25, 50, 75, 90% fell drastically to 100%, 0%, 0% and 0% respectively. Suppl. Figures 11,12 visualize the results from the smaller experiment in the general and hypoxia/normoxia comparison.

#### Community Detection on optimized networks and maximum betweenness centrality per community, provide a gene signature for each mutation type pancancer, in cell lines and tumour samples separately, highlighting the functional differences

We performed community detection Blondel et al. [2008] using the well established *Louvain* method on all optimized networks. This method attempts to create a graph partition so that the modularity metric is maximized. A bigger value in the modularity metric means that the identified communities are more tightly connected as independent hubs. The process is heuristic and stops when the value of the modularity metric can no longer be improved by the means of moving entities (genes) in the network across different hubs. We then disconnected the networks across all detected communities and performed maximum betweenness centrality Li et al. [2017]; Barthélemy [2004] scoring across all hubs. By the union of all the highest scoring genes per community, in terms of the centrality score, we created gene signatures (see Figure 6) for each different mutation type and for all cancer types/sub-types in both CCLE and TCGA separately (Suppl. Material/centrality).

**Figure 6:**
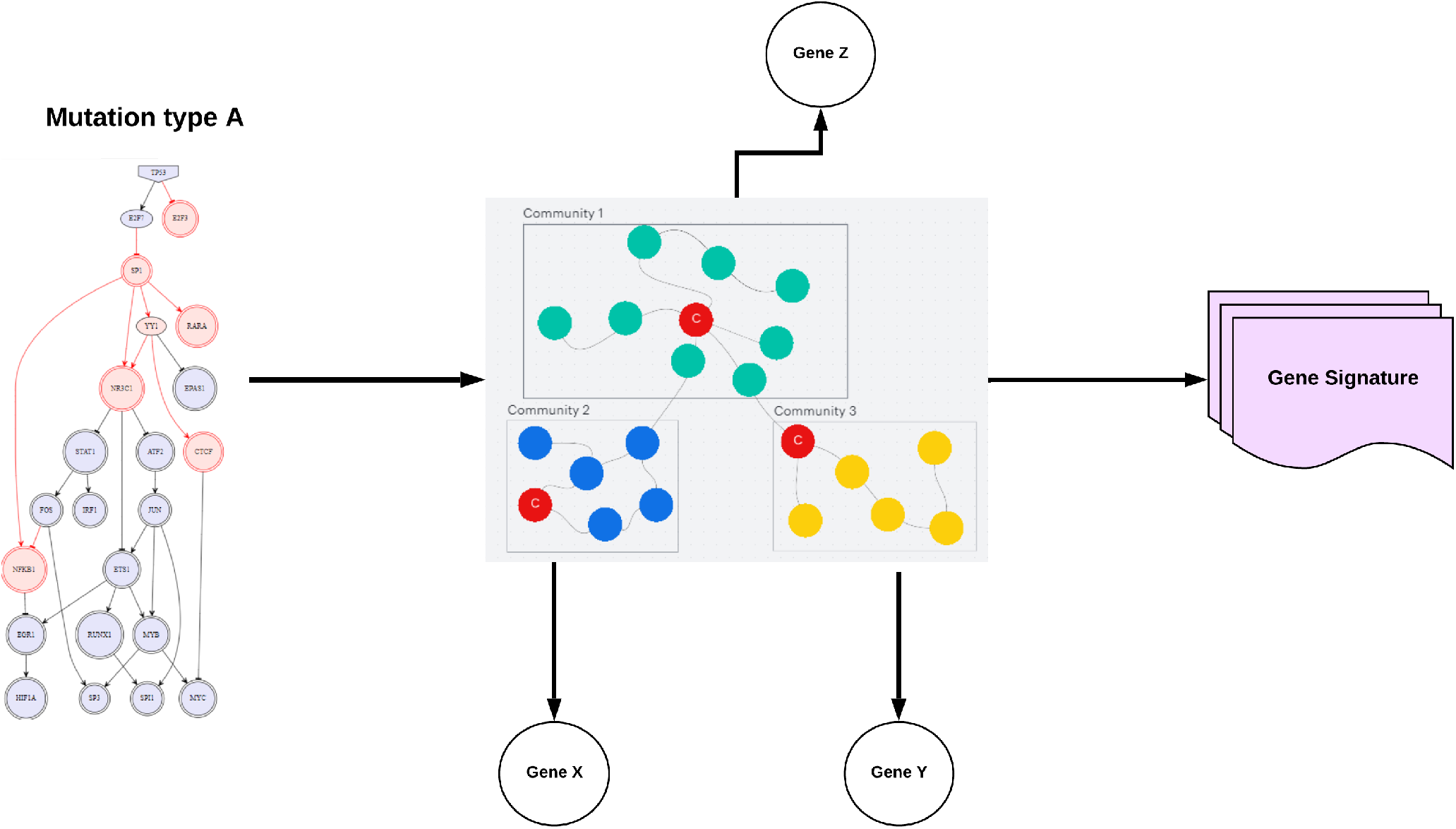
The methodological process to extract gene signatures per mutation from the optimized and reconstructed networks. On the left we see an optimized network which corresponds to a specific mutation type (A) for *TP53*. This network is then partitioned using community detection (Louvain Method) and then each community is mapped to a single gene (as shown, X, Y and Z.) by the maximum betweeness centrality score (red node). These genes are then merged and form the gene signature for the specific network. Then, taking the overlap of all these gene signatures extracted from all networks that correspond to the same mutation type of *TP53*, provide the meta-signature for this mutation type. This process has been repeated across both CCCLE and TCGA data sets.

We then merged the signatures per mutation type across all cancer types/sub-types. Hence, we constructed nine meta-signatures, one for each different mutation type but in a pan-cancer manner for CCLE and TCGA. We present the distinct intersections of the sets of signatures(using *upSetR* Conway et al. [2017]) across all different mutation types in both TCGA (Figure 7(a)) and CCLE (Figure 7(b)). We can see that the missense mutation signature, the most prevalent in terms of frequency of occurrence across different cancer types, shares only one gene with deleterious mutations in TCGA and six with non-Deleterious in CCLE. This indicates the specificity of this meta-signature in comparison to the rest.

**Figure 7:**
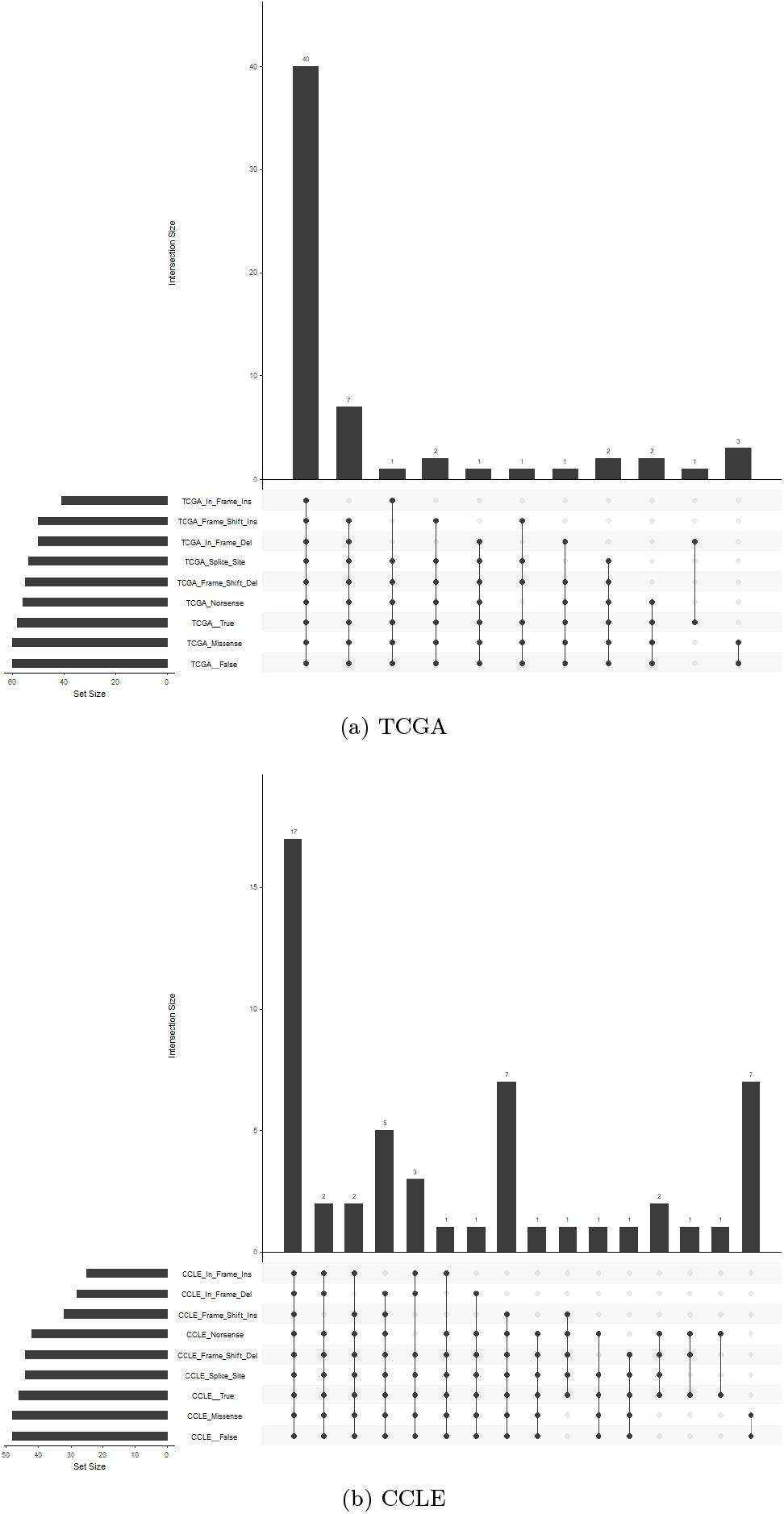
*TP53* mutational meta-signatures (across all cancer types) for TCGA 7(a) and CCLE 7(b) using upSetR Conway et al. [2017] and stratifying by degree. The signatures fluctuate in the number of genes involved approximately from 40 to 60 genes per mutation, in both cell lines and tumour samples. We can see the similarity in the number of genes shared across signatures in both CCLE and TCGA in the first column (all signatures), having 17 common genes in CCLE and 40 common in TCGA. Notably, missense mutations (the most prevalent across cancers) share seven genes in CCLE with non-deleterious signature and three genes with non-deleterious signature in TCGA, highlighting the specificity of the missense signature pan-cancer.

## DISCUSSION

Here, we present a new computational systems network approach, assessing the functional effects of the mutational landscape of the most mutated gene in human cancer, *TP53*, across cell lines and tumour samples. To our knowledge, this approach is new in the sense that it attempts to evaluate whether different mutations impact the regulon and interacting pathways of a transcription factor, in this case *TP53*, in a substantially different way, as opposed to simply differentiating between WT and MT as previous research focused on. Conversely, we are also able to assess whether the same types of mutation cluster across the similarity of the reconstructed networks. Importantly, our approach is not limited to the downstream regulon, but it also accounts for potential upstream network rewiring, which can involve other transcription factors and interacting pathways.

To evaluate our approach we used genomics and transcriptomics data from large public databases, together with previous knowledge of network biology. We evaluated the efficacy of the gene signature from DoRothEA (the *TP53* regulon we used) on its ability to predict the status of these differential features in both CCLE and TCGA versus another well-known *TP53* signature. Although the regulon classifier distinguished better any type of mutation of *TP53* versus the 4-gene classifier, both were unable to predict effectively the type of the most prevalent *TP53* mutation (missense) or the deleterious function of any mutation.

Thus, we resorted to network biology optimization to extract the optimised networks (the regulon of *TP53*) by using perturbation experiments sequentially; each experiment maps to a unique *TP53* mutation and the perturbation depends on the deleterious effect of the variant classification (deleterious: knockdown, non-deleterious: activated). In this way, we were able to calculate precisely the topological similarities of all possible pairs of network comparisons across all *TP53* mutations found in both data-bases. In turn, we classified the similarity strength based on either random pooling (no underlying common mutational feature/total-general similarity in tables and figures) or against same type of mutation, same deleterious function or an identified *hotspot* mutation for *TP53*.

Across different types of cancers, the strength of the signal we infer about the relationship of the mutational status of *TP53* and the status of its regulon downstream was found to fluctuate, with some cancers showing a stronger signal while others remained more invariant. However, the direct conclusion is that when we compare the optimized networks based on same mutation type or deleterious function of the mutation, or same protein change and identified hotspot mutations, we obtain more similar networks as opposed to random comparison across all networks (which account for mutated only samples of TP53). This is consistent across 22 types of cancer in cell lines (CCLE) and 54 sub-types of cancer in tumor samples (TCGA), further implying that our methodological approach can unveil the true phenotypic impact and functional characterization of transcription factors.

Using highly established community detection methods and centrality metrics, we were able to extract gene signatures (a sub-set of the regulon of *TP53*) for all optimized networks. By stratified and unifying per mutation type first and then per cancer type, we were able to construct nine (one per mutation type) meta-signatures, representing the union of all signatures extracted per mutation type across either cell lines (CCLE) or tumour samples (TCGA). The comparison of the signatures clearly demonstrates that although a significant proportion of genes overlaps across all signatures, very few are then shared across all the possible comparative combinations among the signatures. This shows that the regulatory rewiring of the *TP53* regulon has distinct differences per mutation type, once we reconstruct the networks and stratify per mutation type or deleterious function of the mutation for the gene. In essence, the gene signatures provide an indirect way to *visualize* almost twenty thousand of optimized networks.

Mapping the transcriptome of a given sample and therefore potentially an individual cancer patient, to unique single *TP53* mutations and studying the effect on its regulon can pave the wave to personalized treatments, tailored specifically for each case. This is the epitome of personalized medicine and patient stratification in the endeavour of improving prognosis and treatment in cancer patients on severity and disease progression. Finally, the approach can be extended to study more generally genetic variation, as it zooms in on mutation type, searching for mechanistic effects explainable by the rewired networks.

## METHODS

We combined multiple computational techniques and modeling schemes to provide an integrated platform for rapidly performing experiments given CCLE or TCGA data-sets. We extracted 587 genes as the predicted regulon from DoRothEA using Cytoscape. The regulon itself adapts to the optimization output each time; given the initial network of genes and their known interactions, some may be dismissed as nodes or their edge type might be altered to fit the *expression (training*) data, which contain the gene expression profiles (GEP) extracted from the Pan-Cancer Cancer Cell Line Encyclopedia (CCLE) and the Tumor Cancer Genome Atlas (TCGA) data-sets each time. The input data are usually perturbation simulations, where deleterious mutations in *TP53* are assumed to render the protein dysfunctional, allowing the model to predict the regulon using optimization to best fit the gene expression data. The inference achieved in this way can serve as a basis to create a network that allows us to study the downstream effects of specific p53 mutations. We used the full transcriptome as an expression input matrix, to make sure we become fully inclusive in interactions that might help reconstruct the networks more accurately.

Once the signature and the expression matrix are selected, formatted and finalized, CARNIVAL attempts to optimize the selected PKN network by using information from the RNAseq expression values, the PKN and the measurements for the transcription factors included in the initial network (Normalized Enrichment Scores as those extracted from the R package VIPER).

The resulting optimized network assigns states (node-wise) and relationships (edge-wise). More specifically, the TF will be marked as the perturbation node, and downstream of this we will see the nodes (genes/proteins) being either up-regulated (red) or down-regulated (blue) with the corresponding edges acting either as an activator or inhibitor, from source to target (the PKN is directed).

Each optimized network (assuming the optimizer converged) is stored and characterized by the single *TP53* mutation it corresponds to. We then computationally compare all networks across each other (the number of possible comparisons then reduces to the basic principles of counting; as we wish to avoid the main diagonal and our matrix of comparisons is symmetric above and below; order does not count. Thus we derive the number of comparisons required as the combination of choosing 2 from N, 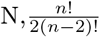 ^2^).

Comparisons are being calculated on the basis of four different metrics: i) general, meaning that we do not filter the compared networks across any feature, so this amounts to the max number of possible comparisons, ii) we first filter the networks across the same type of variant classification (type of mutation), iii) we use the deleterious attribute of *TP53* in each condition/mutation/optimized network, and finally iv) we filter across only mutations that have been characterized as known hotspots for *TP53*. Each metric besides the first one shrinks the number of possible combinations as expected. However, the results are phrased in percentages to reflect generality and not resolution. We used four different percentages of similarity to compare all four different metrics/settings: (25%, 50%, 75% and 90%).

### Cancer Cell Line Encyclopedia (CCLE)

We used the Cancer Cell Line Encyclopedia omic data-sets Ghandi et al. [2019], *DepMap Public to train the networks*^3^ available at https://depmap.org/portal/download/. We collected all the cell lines corresponding to each of the 22 different cancer types in a .csv file and used it as an input for our analysis tool. The expression and mutation profiles from DepMap and the cell lines corresponding to each cancer type comprise the total input to our CCLE analysis tool. Specifically, we downloaded the.csv files for expression (RNAseq [*log*2 + 1]) and the mutational profiles for all genes. The expression profiles serve as an input to both the linear regression and the CARNIVAL optimization modules. The mutational profiles are used for CARNIVAL to extract mutational status and further features (such as deleterious function) for *TP53*, which is the perturbed input in the network for all optimizations performed.

### The Cancer Genome Atlas (TCGA)

We used the survival data as provided on the NIH National Cancer Institute Genomic Data Commons website at http://api.gdc.cancer.gov/data/3586c0da-64d0-4b74-a449-5ff4d9136611 and https://api.gdc.cancer.gov/data/1b5f413e-a8d1-4d10-92eb-7c4ae739ed81, respectively. We then combined the expression profiles with mutations downloaded from the *cBioPortal* (https://www.cbioportal.org/). This provides the full input for the pipeline of the developed platform to perform either Generalized Linear Modelling (GLM) or CARNIVAL experiments. We extract the same semantic fields such as deleterious function, as initially described for the CCLE data-sets in a seamlessly equivalent way.

### Radiation Experiment

H460 (p53 wild-type Non-small cell lung cancer) cells were grown in parallel in two 175 cm2 flasks with RPMI-1640 media supplemented with Fetal Bovine Serum (10%), Penicillin-Streptomycin (1%) and L-glutamine (1%). Media was changed three times per week and passaged with trypsin (1%) on approaching confluency using aseptic technique. A cobalt gamma irradiator was used to deliver 50/60Gy in 2Gy per fraction over 5/6 weeks. Cell lines were authenticated via genomic analysis (Northgene) and underwent regular mycoplasma testing. Following irradiation sub-lines were confirmed to be radio-resistant versus parental via clonogenic assay. An RNAseq experiment was performed for the unirradiated parental (H460 BASE) and sublines (H460-60A and H460-50B) at 5 time points; pre-(0hrs) and post-(2, 6, 12 and 24hrs) a further 2Gy irradiation. Cells were seeded to 6 well plates the prior day and harvested per timepoint, three biological replicates were performed per condition and total RNA extracted using TriZol (Thermofisher) following manufacturer’s protocol. Library preparation was performed with Lexogen Quantseq 3’ RNA kit and sequencing with the Illumina NovaSeq platform by Welcome Trust Centre for Human Genetics Oxford (WTCHG).

### Hypoxid Experiment

#### Cell culture

All cell lines used (MCF7 Cat HTB-22 RRID:CVCL_0031; MDA-MB-231 Cat CRM-HTB-26 RRID:CVCL_0062; MDA-MB-453 Cat HTB-131 RRID:CVCL_0418; MDA-MB-468 Cat HTB-132 RRID:CVCL_0419; and HCC1806 CatCRL-2335 RRID:CVCL_1258) were purchased from ATCC. They were routinely cultured in DMEM low glucose (1g/l) and supplemented with 10% FBS no longer than 20 passages They were mycoplasma tested every 3 months and authenticated during the course of this project.

For the larger experiment, cells were cultured in different glucose (Gluc) and glutamine (Gln) levels as follows: medium A) 1mM Gln, 5mM Gluc; medium C) 4mM Gln, 5mM Gluc; medium D) 1mM Gln, 2mM Gluc; and medium F) 4mM Gln, 2mM Gluc; and subjected to normoxia (21%), 1% or 0.1% hypoxia for 24h and 48h using an InVivO2 chamber (Baker). For the smaller experiment, cells were cultured in either: medium C, DMEM high glucose (4.5g/L) 4mM Gln, Human Plasma Like Medium (HPLM; Gibco A4899101), or Plasmax Vande Voorde et al. [2019]. All media were supplemented with 10% FBS. Cells were seeded the day prior to the experiment, then cultured under normoxia (21%) or hypoxia (1%) for 24 hours.

#### RNA extraction and DNAse treatment

RNA from each experiment was extracted using the mirVana™miRNA Isolation Kit (AM1560, Thermo Fisher Scientific) and DNAse treated with TURBO DNA-free™Kit (AM1907) following the manufacturer’s instructions.

### RENOIR - Generalized Linear Models

REsampliNg methOds for machIne leaRning (RENOIR) is an *R* package developed to obtain reliable estimates of the true error of a trained model, i.e. the difference between the true value and the approximation resulting from the model prediction. The estimate of the true error is important as it allows us to understand how well the developed model generalised to unseen data. RENOIR does this by using a multiple sampling strategy that can be summarised in the pseudo-code below:

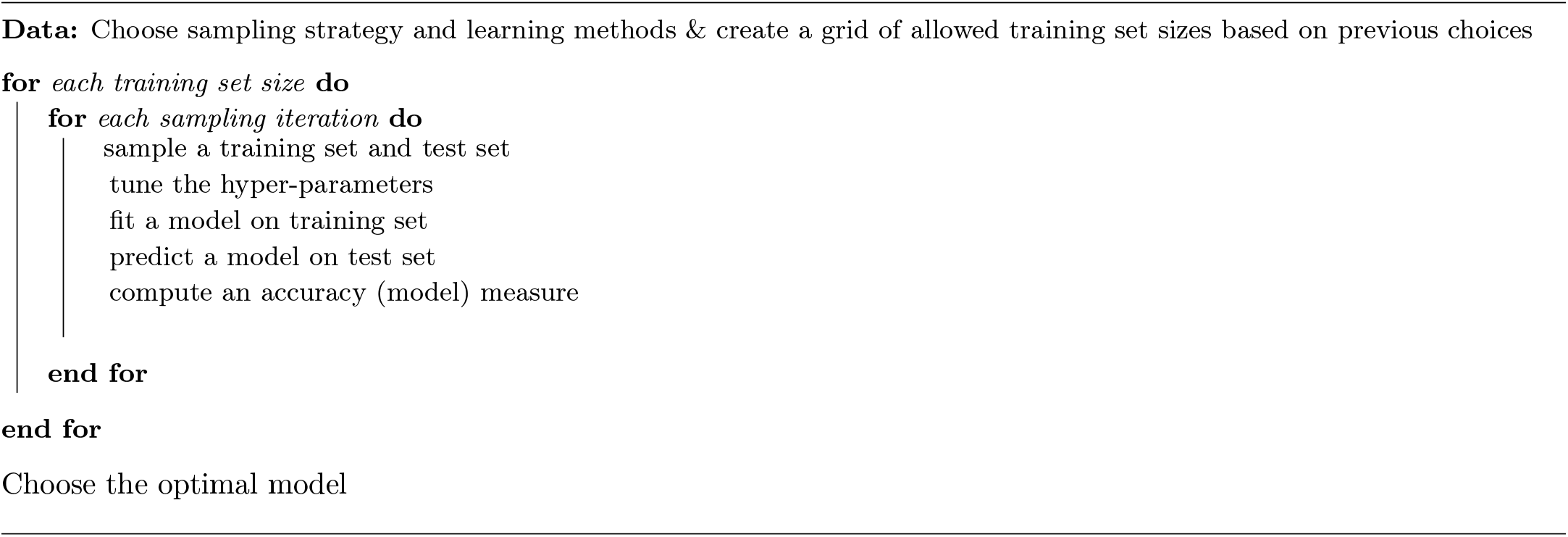

The sampling strategy used in this work is a multiple random sampling, while the used learning method is GLM via elastic net regularisation. The mixing hyper-parameter alpha was tuned via 10-fold cross-validated grid search. A feature screening was performed inside cross-validation to reduce the high-dimensionality of the feature space and select the variables most strongly related with the response. The package has been developed by Dr Alessandro Barberis and Prof Francesca M. Buffa at the department of Oncology, University of Oxford, UK.

### CARNIVAL - Mathematical Programming optimization

CARNIVAL Liu et al. [2019] uses Mixed-Integer Linear Programming to optimize/train a given Prior Knowledge Network (PKN) which serves as a starting point on how the topology of the interactions is delineated. For a given single condition, the model then fits the experimental data (expression) for the nodes of the network altering either the type of the interaction or dropping a node (gene-protein) as the size of the network is penalized given the Transcription Factors (included in the PKN) activities. The resulting optimized network minimizes the mismatch between the measurements and the predicted states (up or down regulation) of the network nodes. In this way, a single mutation of *TP53* mapped as a unique condition and a perturbation input for our network can be assessed for its downstream impact on the regulon (all the down stream identified targets of *TP53* using DoRothEA), by comparing the two networks. Effectively, we then stratify the comparison across the same settings we categorized earlier the binomial classifiers: i) mutation type, ii) deleterious function and iii) hotspots. We then analyze how strong of a signal we receive for low, medium and highly similar networks (correspondingly 50, 75 and above 90% similarity scores) to assess which of the settings analyzed explains a similar downstream effect for each *TP53* mutation.

### Network comparison

CARNIVAL outputs the .dot files of the optimized networks which we then first read with the *R* package *sna* and then convert to *igraph* objects so we can apply the intersection of the two graphs. Let *G_A_*, *G_B_* be the two to-be-compared optimized gene networks as those output from CARNIVAL. Then let *G_A_* = {*E_A_, V_A_*} and *G_B_* = {*E_B_, V_B_* } the sets of edges and vertices correspondingly for the two graphs. The intersection of the two networks is calculated as the union of all edges present in both graphs. More formally, if *G* is the intersection of *G_A_, G_B_*:

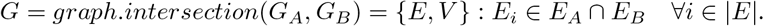

Assume now without loss of generality that |*G_A_*| ≥ |*G_B_*|, where |*G*| is the size (number of edges) of graph *G*. Then, a score of similarity *S_AB_* can be calculated between *G_A_*, *G_B_* as follows:

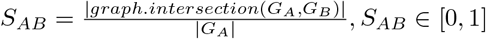

This intersection takes into account node-to-node both direction and type of interaction, therefore yielding a good estimate of how close (or far apart) semantically the compared networks are. The idea of comparing the resulting graphs for similarity we introduce here has also been used in a similar way in the DCI algorithm Belyaeva et al. [2021], where edges that appeared/ disappeared or changed weight are assessed for inference in contrasting two different conditions from the trained respective networks. Network similarity measures have also been used in disease-gene association studies before Sun et al. [2014]. To compare the networks computationally, we create a matrix with each network as a row and column and then take all possible different comparisons. We then exploit the structure of the matrix (symmetric and skipping the main diagonal) and efficiently compute all similarity scores. This procedure is systematically done across all optimized generated networks in both CCLE and TCGA data-sets. An example is shown in figure 3 where we compare two different networks as a downstream effect of different perturbations on *TP53*. Each network is characterized by a unique identifier, as a collection of features as shown at the top of the figure. We merge information about the variant classification, disease type, codon and protein change as well as the deleterious or not effect of the mutation on the gene as a binary (true or false) flag at the end of the identifier string. This enables us to accurately associate any observations to their exact origin and further our understanding of the possible causality.

### Optimization model

The CARNIVAL optimization model is a Mixed-Integer Linear Programming (MILP) model. It is a special case of constrained optimization where we try to optimize (minimize or maximize) a linear function over a set of also linear constraints under the extra limitation that some of our variables have to be reals or integers (or just binary *x* ∈ {0,1}). This belongs to the general family of Linear Programming optimization models, for which deterministic optimal algorithms exist. For this purpose we used the IBM CPLEX solver.

The objective function attempts to predict the status of the optimized nodes in the network as those up-regulated (1) or down-regulated (−1) which has to abide by the so-called *consistency rules* which enforce biologically reasonable interactions between the genes. This becomes an optimization problem as we force to minimize the function over a feasible set of constraints. For further details on the semantics, derivation, and explanation of the mathematical constraints in detail the reader can refer to Liu et al. [2019]. For clarity, the model is also formally given below.

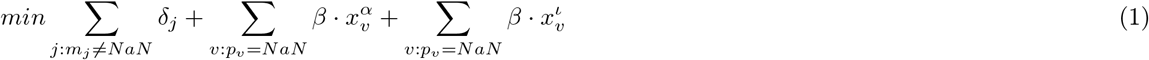

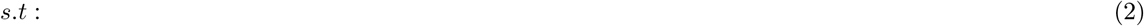

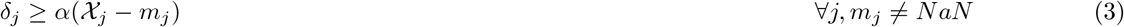

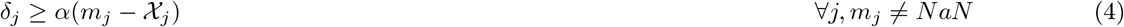

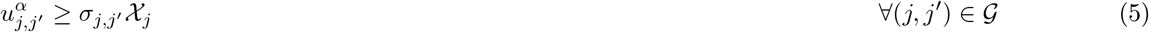

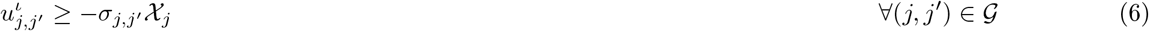

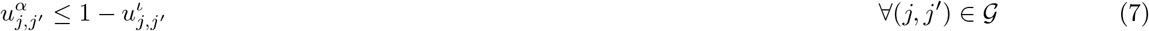

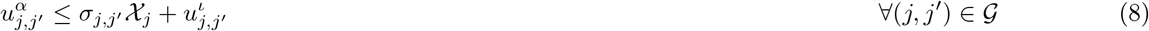

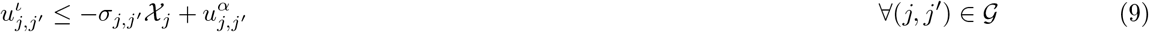

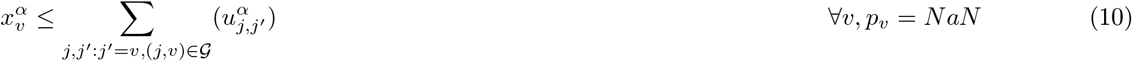

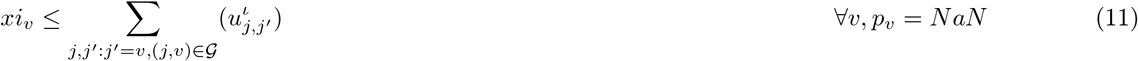

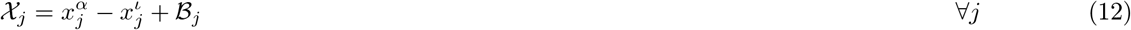

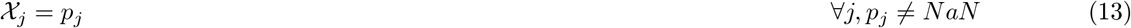

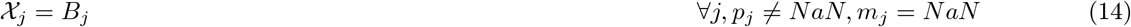

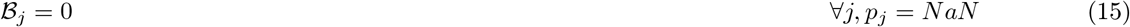

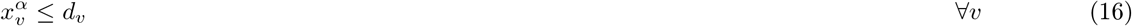

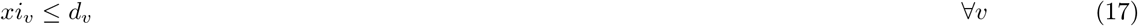

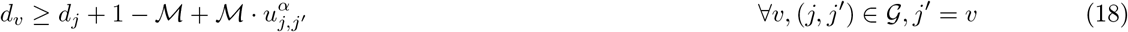

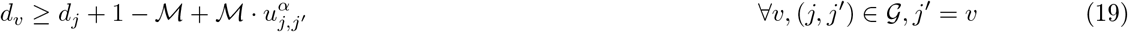

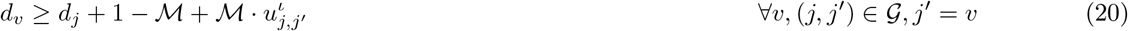

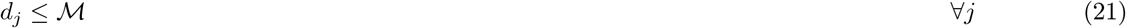

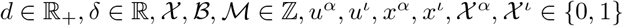

Here, 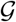 is the original PKN as a graph, {*j,j*’} are edge and alias correspondingly, [*u^α^,u^ι^*] are the potential of an edge to up-regulate or down-regulate a node/gene, [*x^α^,x^ι^*] the potential of a node to be up-regulated/down-regulated, 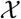 the predicted from the model state of a gene/node, 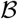 a pseudo-variable for the nodes that are neither measured nor perturbed, *δ* the absolute difference between predictions and measurements and d as the distance measured between nodes; 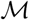 is the upper bound for distance to avoid loops, *β* is a penalty parameter selected by the developers of the model and calibrated appropriately across multiple testing.

### Data and Code Availability

The implementation and the input data with the full results are available as a Shiny app developed in *R* at https://data.mendeley.com/datasets/rn96hp5kw4/2.

## ACKNOWLEDGMENTS

C.P.T., A.M.C. were funded by a European Research Council (ERC) Consolidator Award to F.M.B. (MICROC:772970). A.B. was funded by Cancer Research UK program to F.M.B. P.C. was funded by a Cancer Research UK fellowship. E.G. received funding from J.R.C. for Computational Biomedicine. J.S.R. is thankful to GSK, Sanofi, Travere Therapeutics, Astex Therapeutics for support. The authors also wish to thank Professor Xin Lu for useful discussions which helped improve the quality of the manuscript.

## AUTHOR CONTRIBUTIONS

Conceptualization: C.P.T., F.M.B.; Network Methodology: C.P.T., F.M.B.; Machine Learning Methodology: A.B., F.M.B; Community Detection Methodology: C.P.T.; Formal Analysis: C.P.T., E.G., J.S.R., A.B, F.M.B.; Experiments: A.M.C, P.C, F.H. (supervision F.M.B.); Interpretation: L.V.B., C.P.T., E.G., J.S.R., A.B, F.M.B.; Writing: C.P.T., L.V.B, F.M.B, A.B, Original Draft: C.P.T., L.V.B,F.M.B, A.B; Review & Editing: F.M.B.,C.P.T.,L.V.B,E.G.,J.S.R.

## Supplemental Information

**Suppl. Figure 1:**
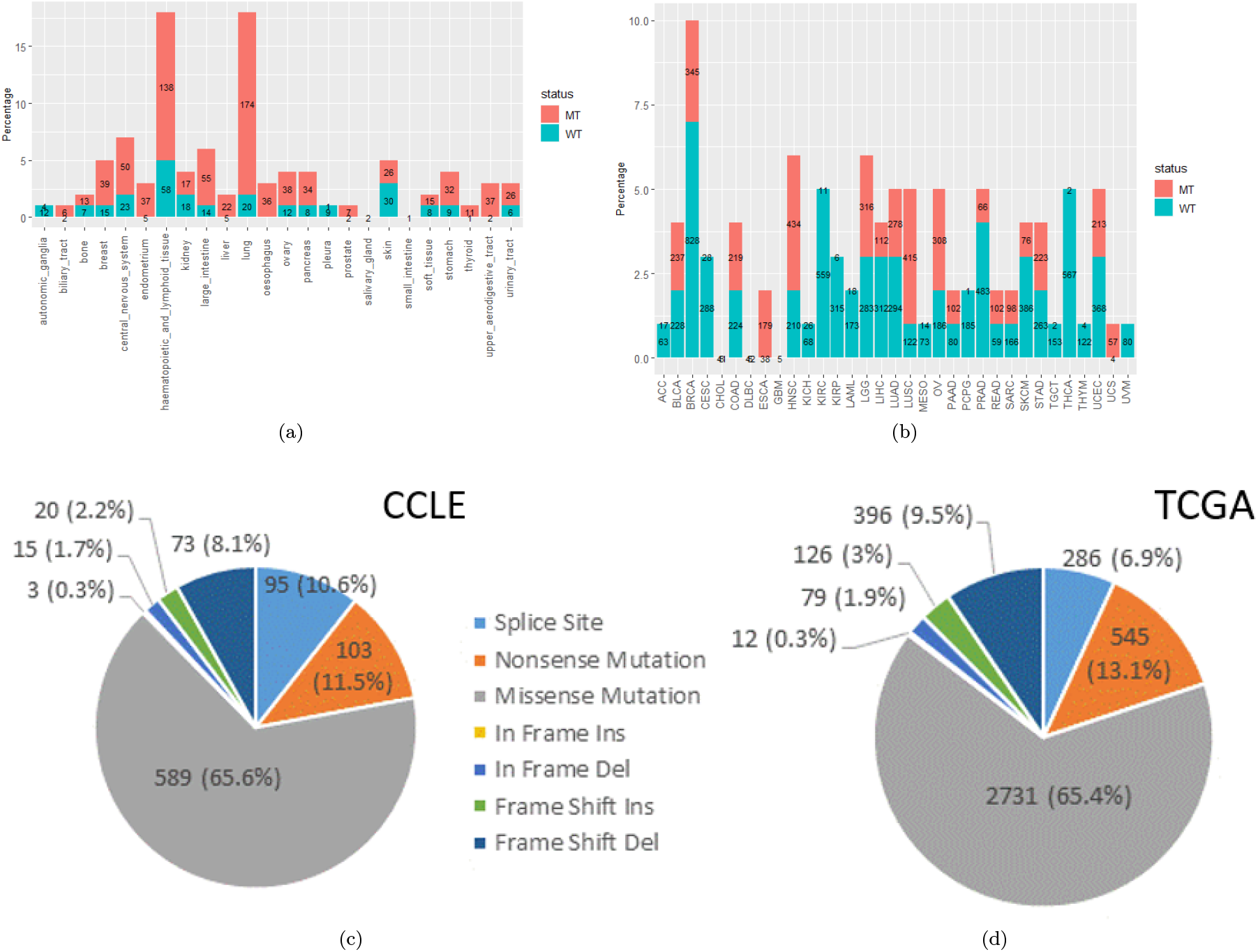
Mutational frequency of TP53 per cancer in CCLE (A) and TCGA (B) (source: https://www.cbioportal.org/). C) TP53 mutational variation count in DepMap CCLE v.20Q2. A total of 898 cell line samples with a TP53 mutation were identified. Ten cell lines had more than one TP53 mutation. D) TP53 mutational variation count in TCGA. A total of 4250 tumour samples with a TP53 mutation were identified.

**Suppl. Figure 2:**
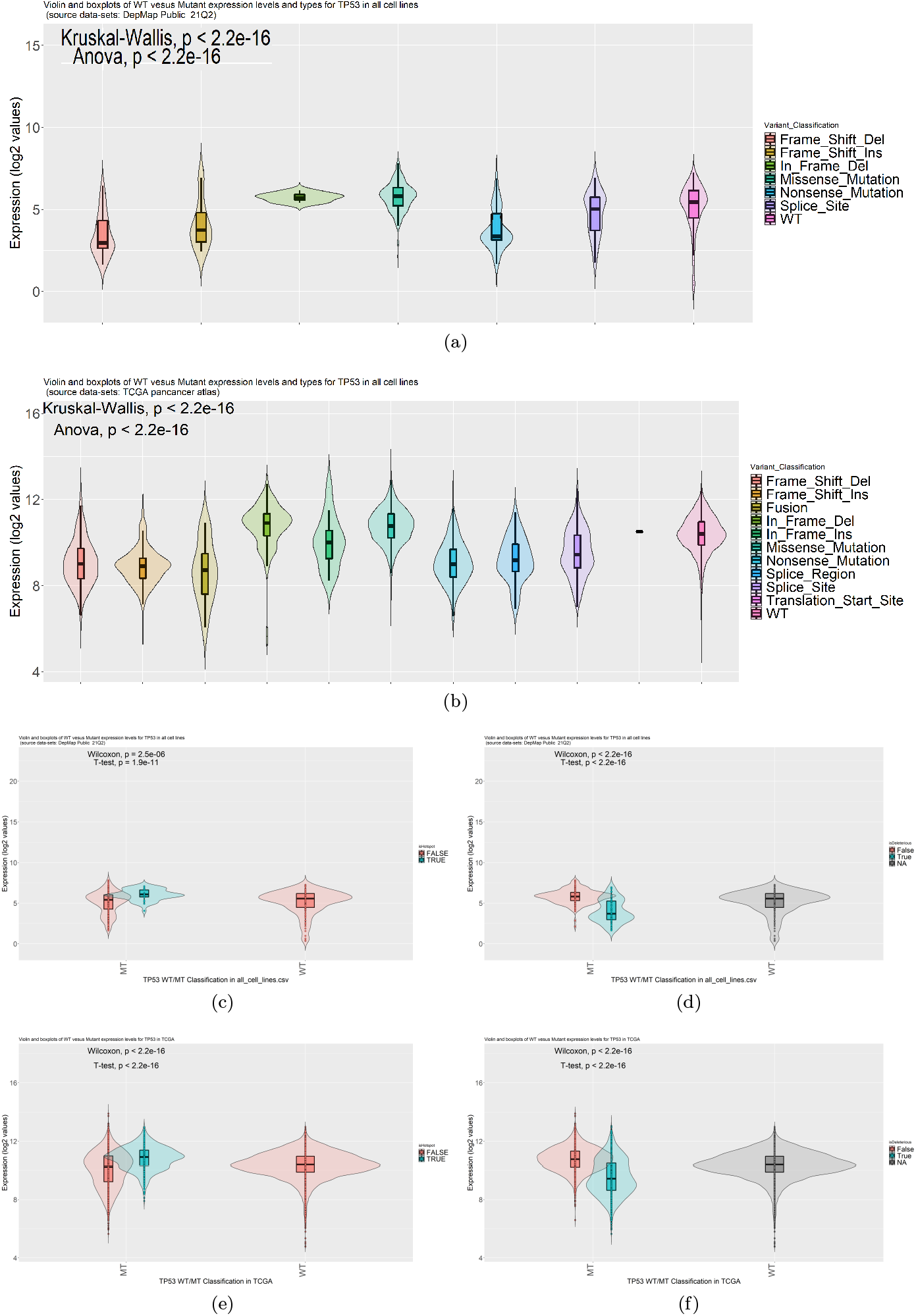
Association between mutations and expression (RNAseq) of TP53 in CCLE and TCGA data. Sources: TCGA PanCancer Atlas and DepMap Public 21Q2. A) Mutation type, B) Hot Spots versus non hot spot mutations, C) deleterious versus non-deleterious mutations as defined on the respective databases. Anova and two-sample tests are shown. Mutation and deleterious annotation was from CCLE and TCGA databases, hotspots were as follows: p.R175H, p.R248Q, p.R273H, p.R248W, p.R273C, p.R282W, p.G245S as defined by Baugh et al. [2017].

**Suppl. Figure 3:**
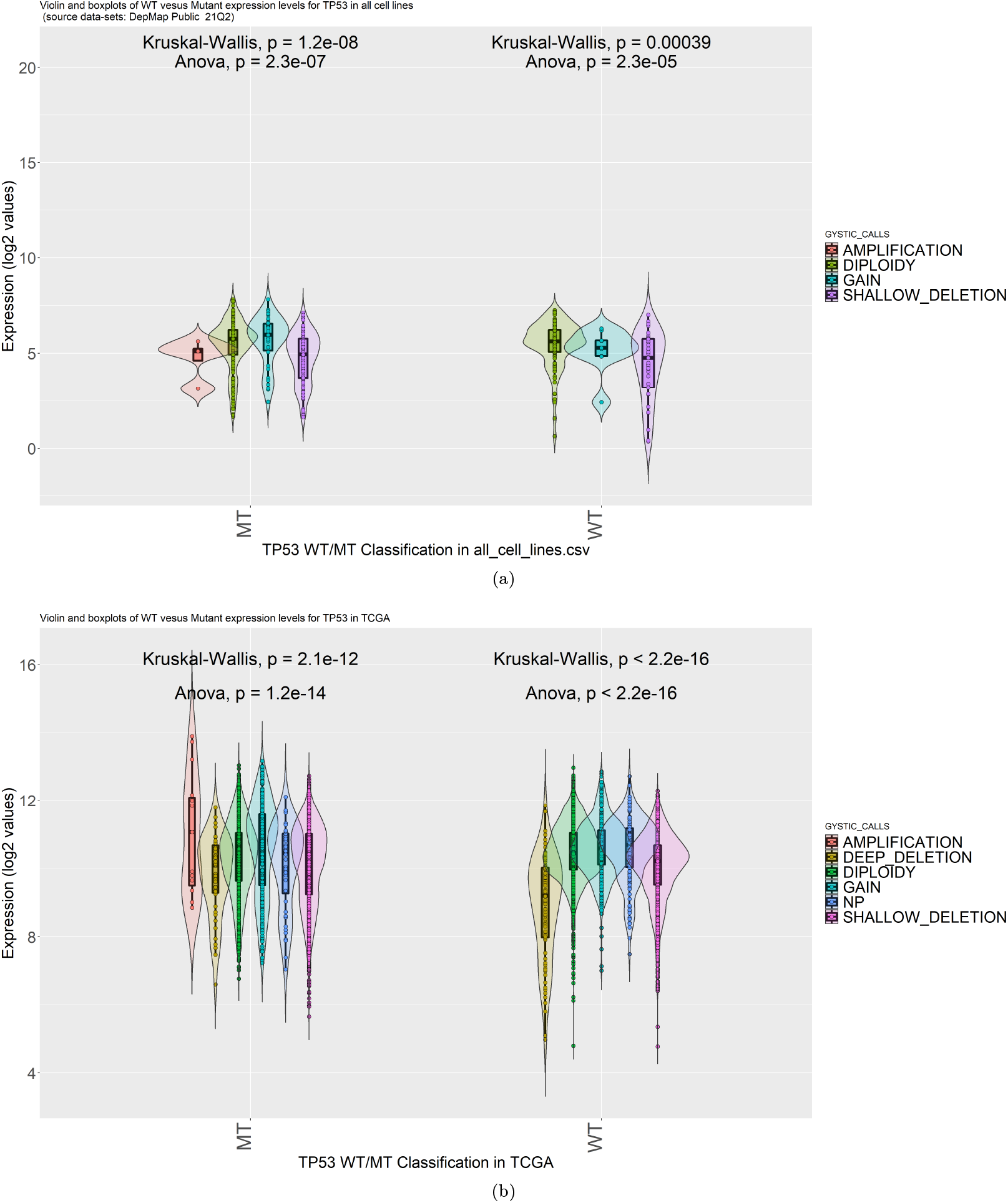
Expression (*log_2_* normalized - see Methods) as a function of TP53 copy number variation, as classified using GYSTIC calls in both CCLE and TCGA.

**Suppl. Figure 4:**
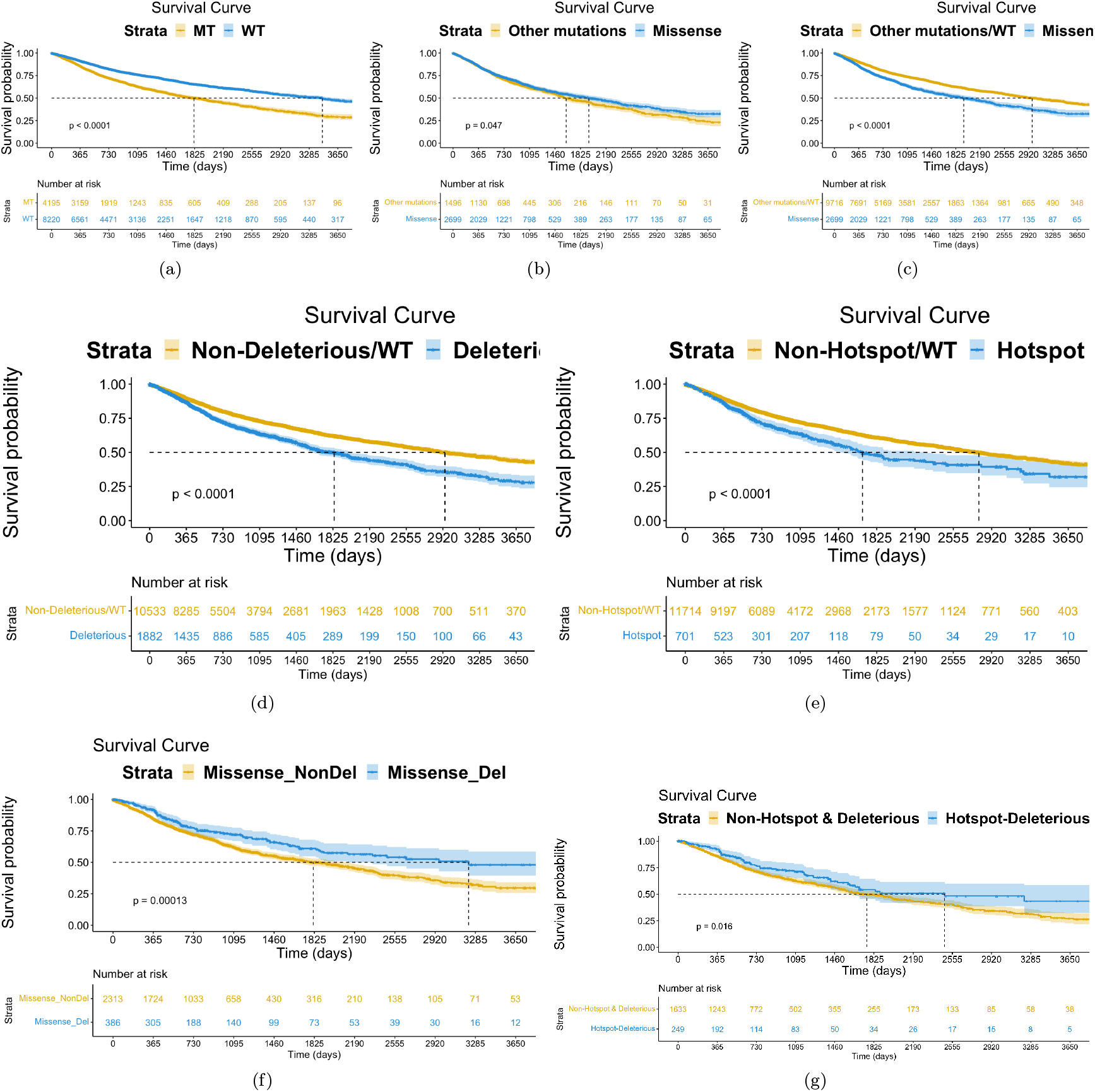
Using the TCGA survival data to plot survival curves on various binomial settings of *TP53* status. The figures show that certain features such as missense mutations affect survival probabilities significantly.

**Suppl. Figure 5:**
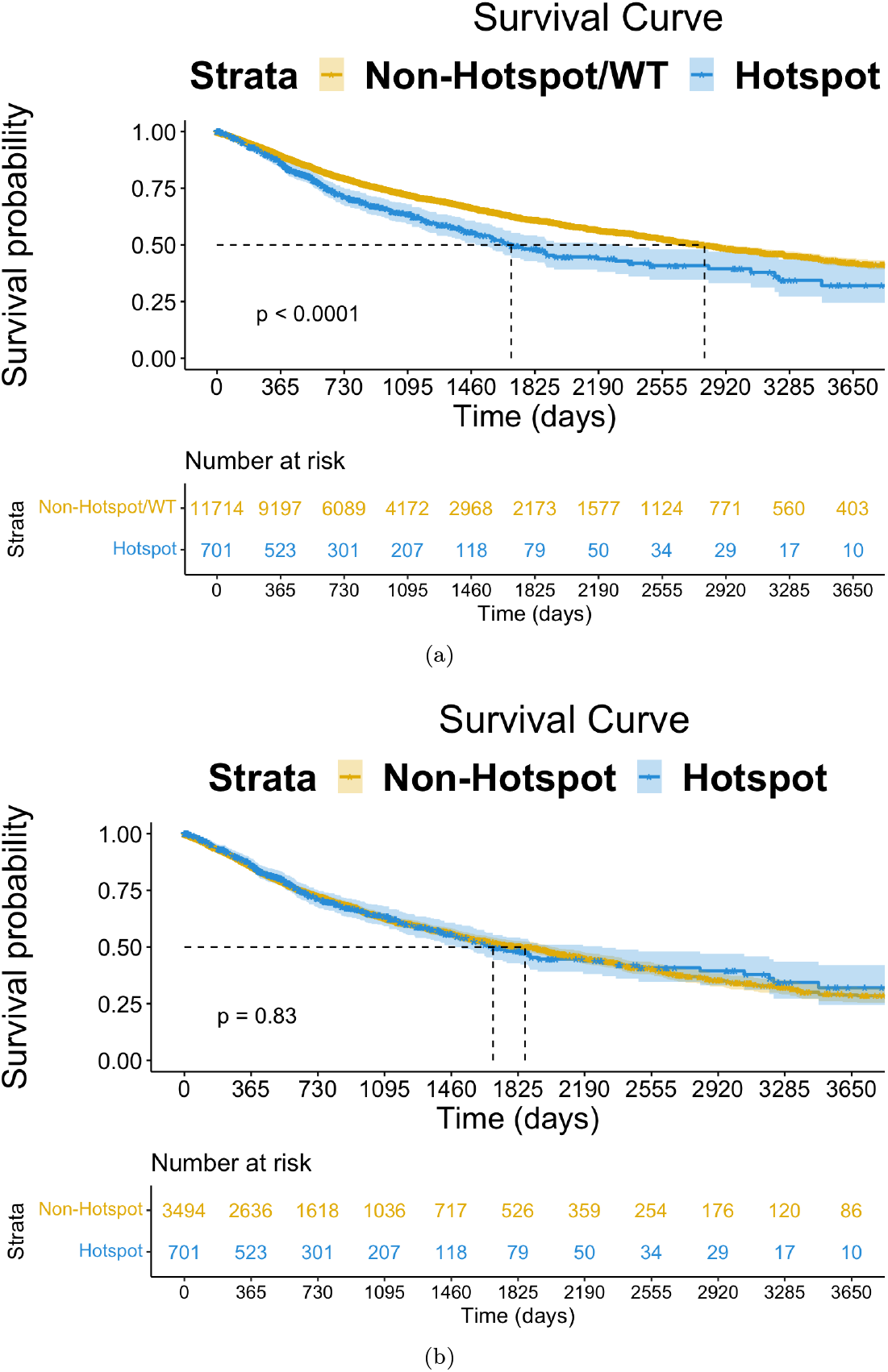
Survival curves in TCGA data across two settings: 5(a) Hotspot mutations of TP53 versus every other mutation (including WT samples) and 5(b) Hotspot mutations of TP53 versus every other mutation (not including WT samples). We can see that we get a statistically significant (p < 0.05) difference only when including the WT samples.

**Suppl. Figure 6:**
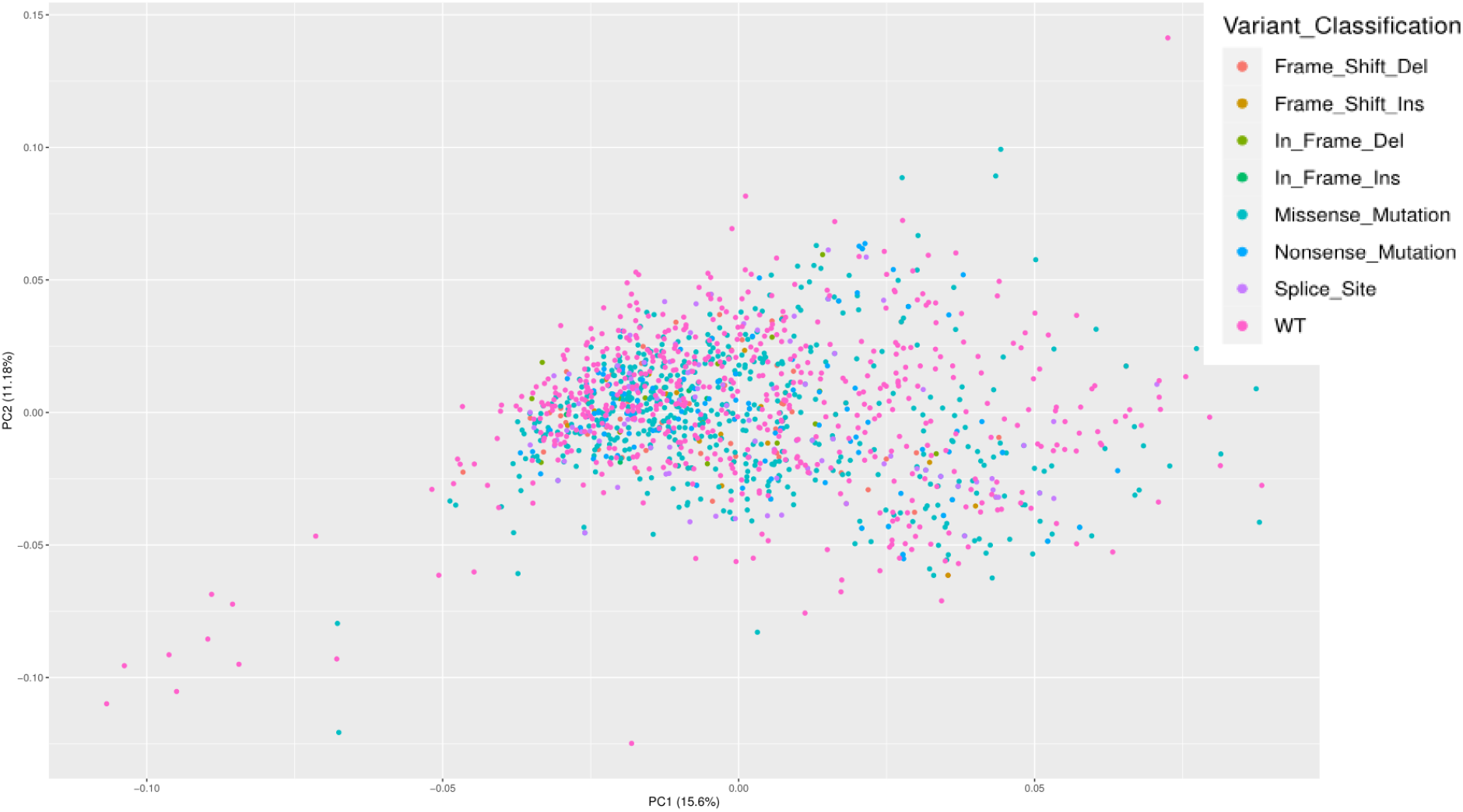
Principal component analysis (PCA) of the expression (RNAseq) of the regulon of TP53 in cell lines samples (CCLE). Principal component 1 (PC1) and 2 (PC2) are shown with the associated variability.

**Suppl. Figure 7:**
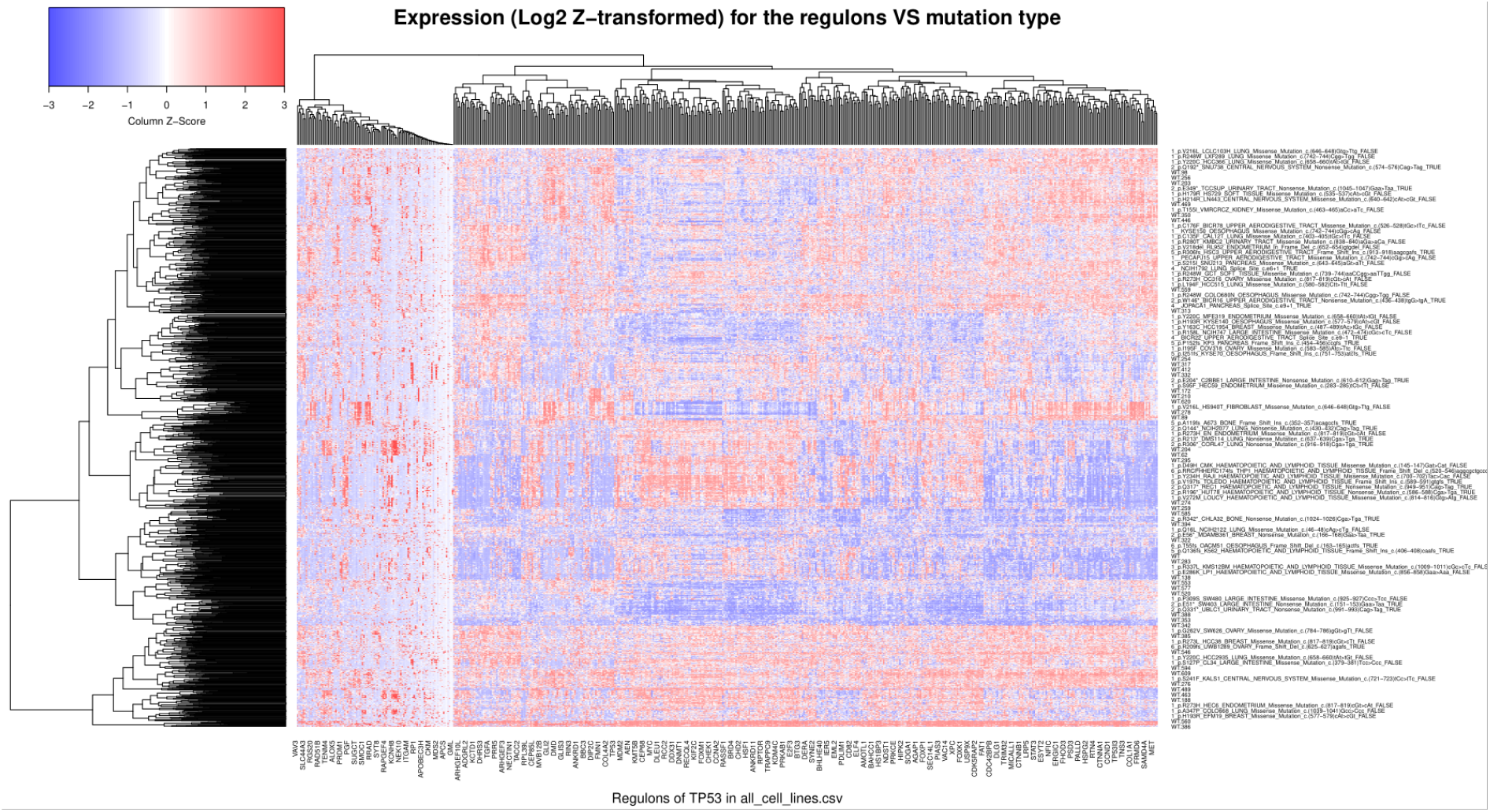
Heat-map of expression (log2) in CCLE for TP53 across all cell lines versus different mutation types and WT samples. The short names on the rows correspond to WT samples, which we would expect to cluster uniformly across the samples, which happens frequently.

**Suppl. Figure 8:**
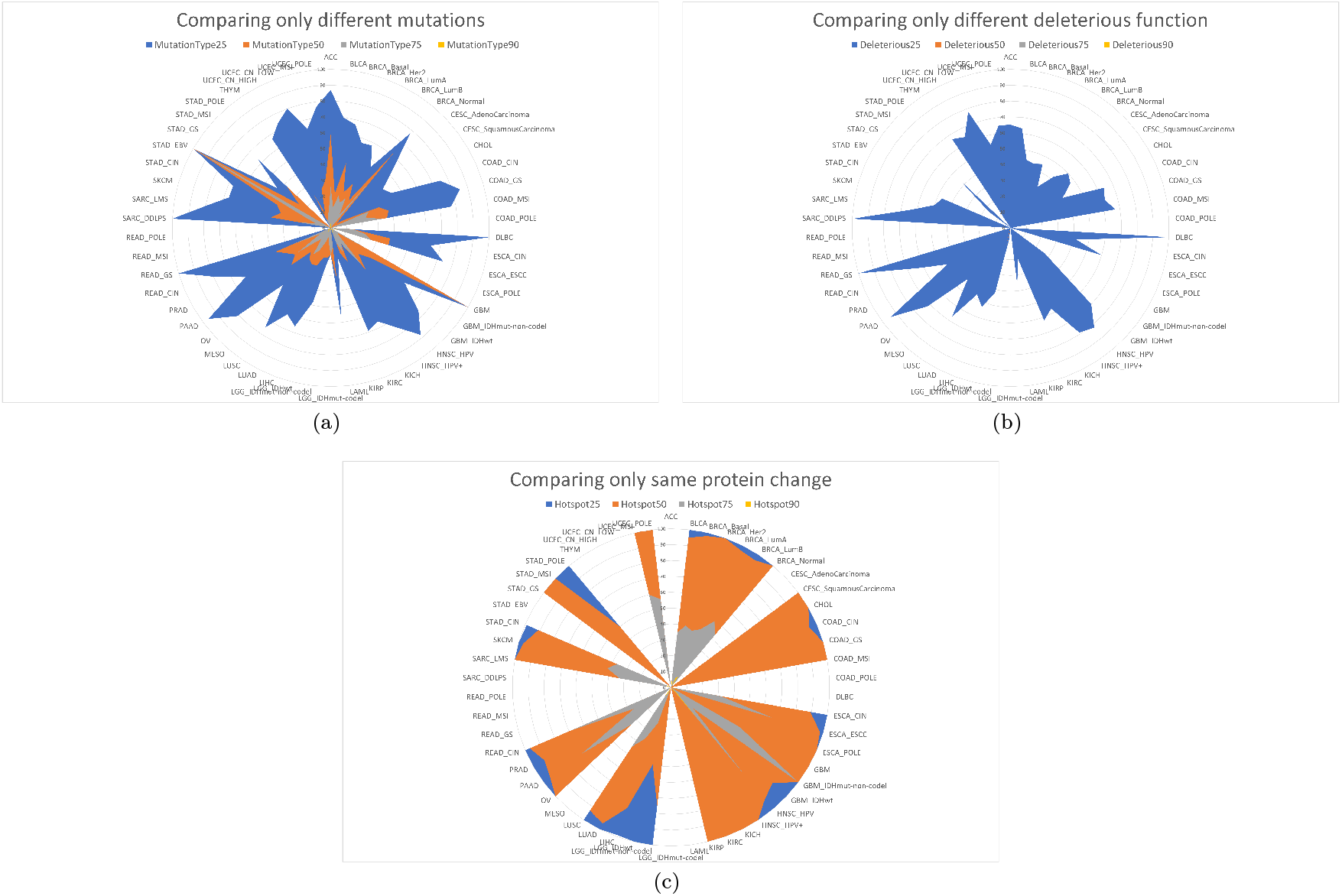
A different aspect of the computational study presented in this figure; we stratify across i) different mutation types, ii) different deleterious function (deleterious and non-deleterious compared only) and lastly iii) exactly same protein change (scarce in occurrence and thus results might not be that rich). We observe how similarity diminishes in the first two cases, as expected from the previous figure.

**Suppl. Figure 9:**
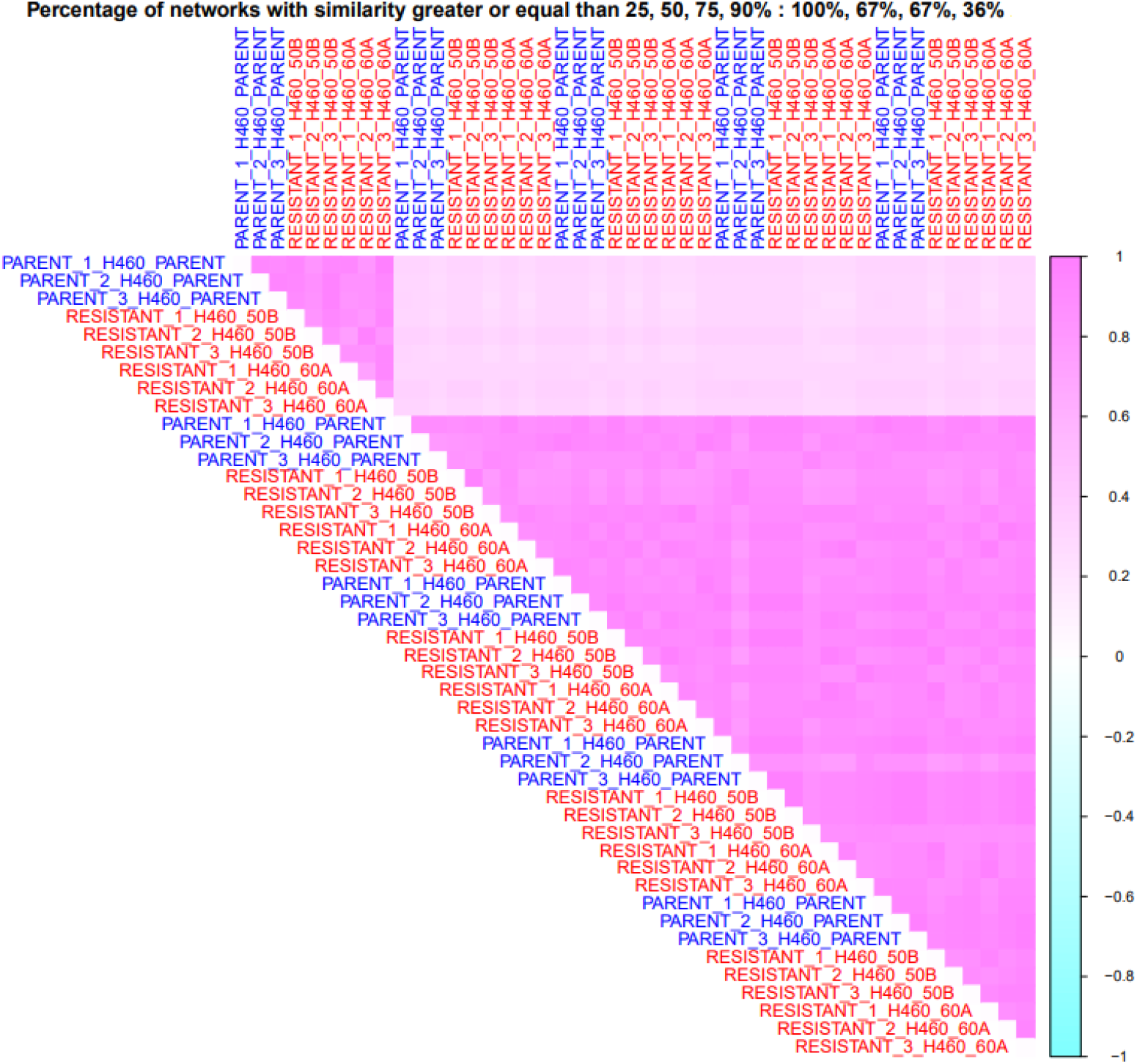
Computationally comparing the optimized networks downstream of *TP53* with and without the application of radiation. The correlation plot shows how similar are the networks in the general case (where even 22% of the networks exhibit more than 90% similarity).

**Suppl. Figure 10:**
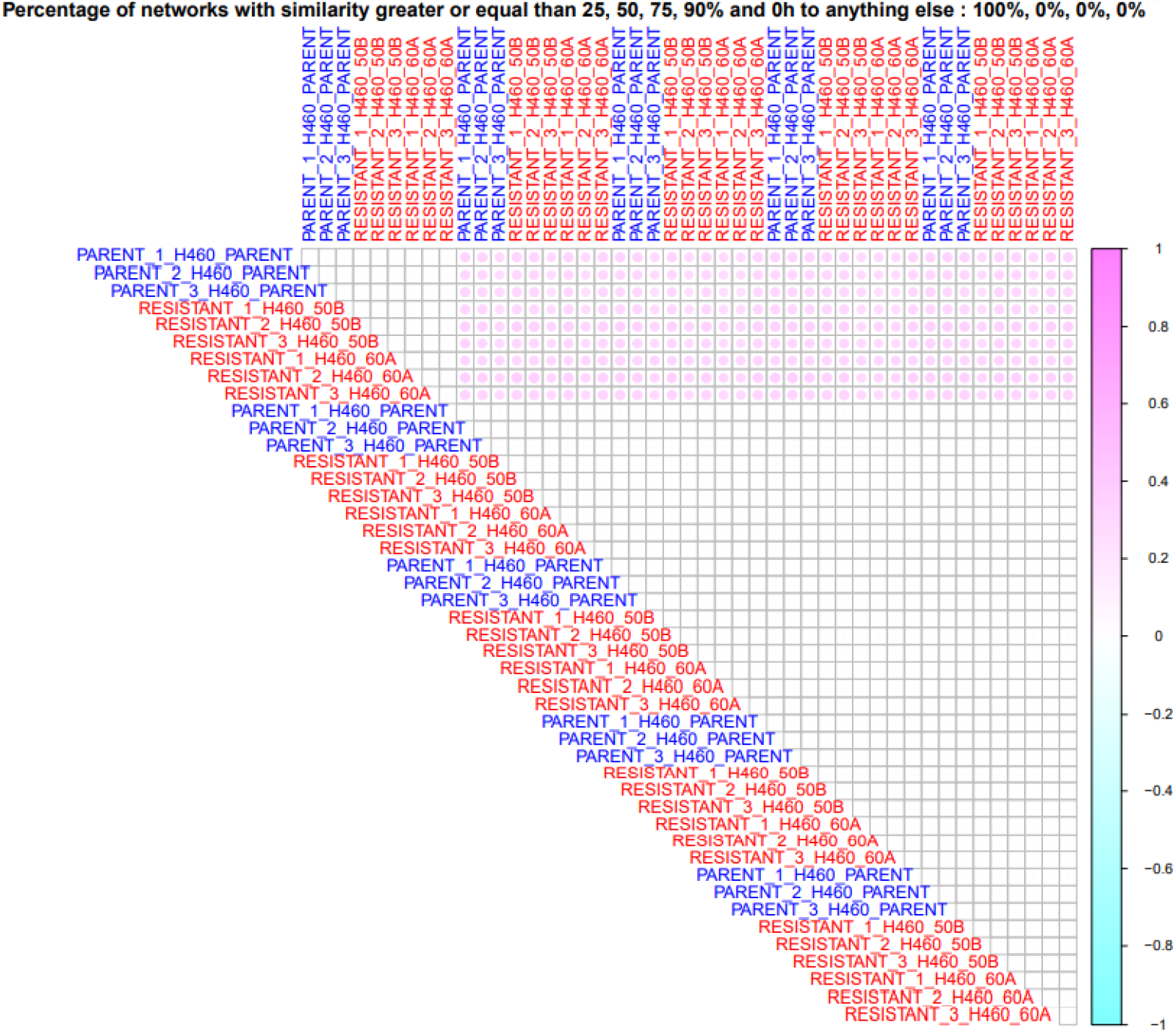
When we compare only between 0h versus everything else (after irradiation), no networks exhibit similarity percentage greater than 25%. More details on the experiment are described in *Methods* section.

**Suppl. Figure 11:**
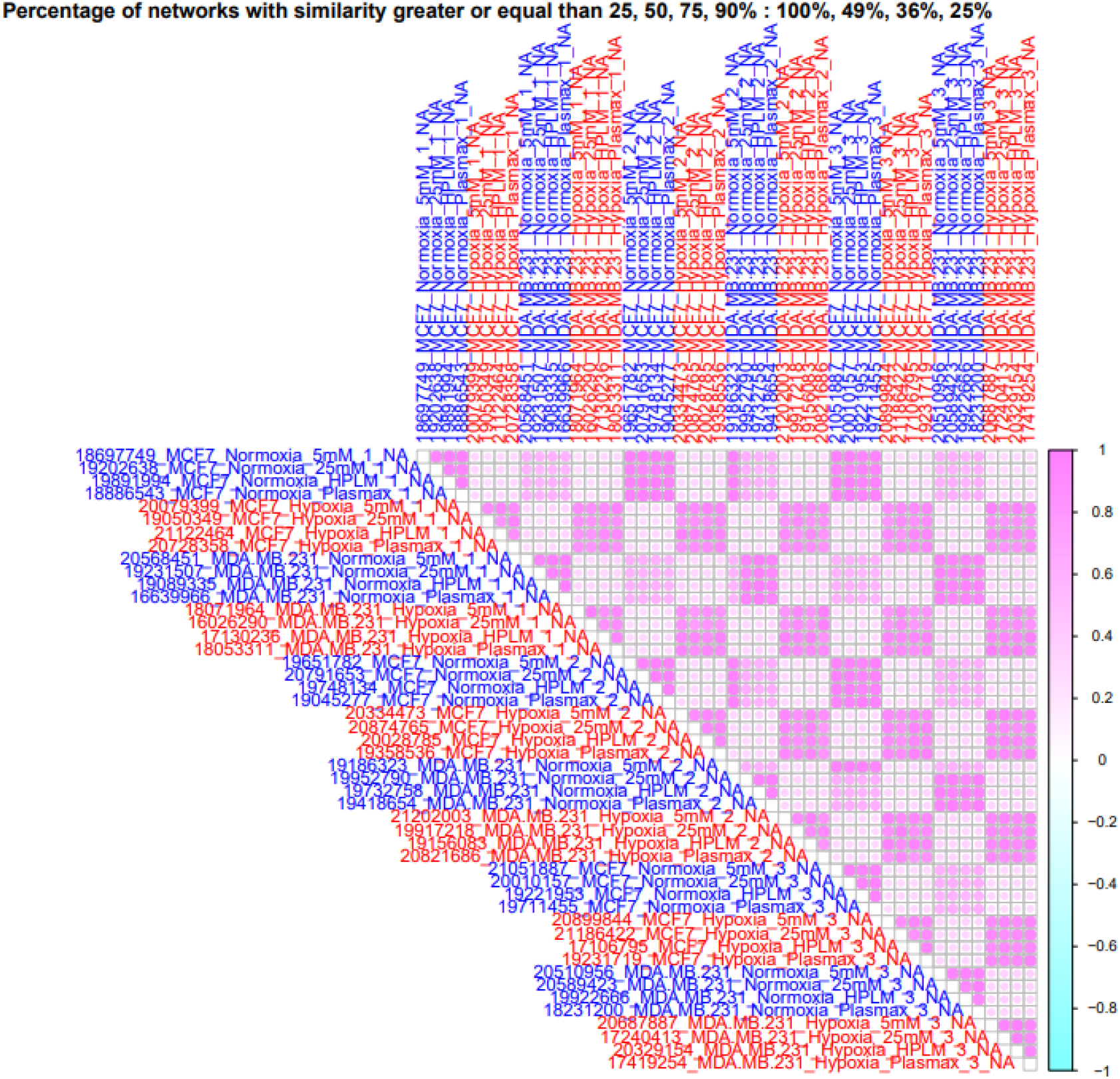
Comparison between all samples. It is evident that similarity increases only when the compared samples come from the same tested conditions, that is, either only hypoxic or normoxic.

**Suppl. Figure 12:**
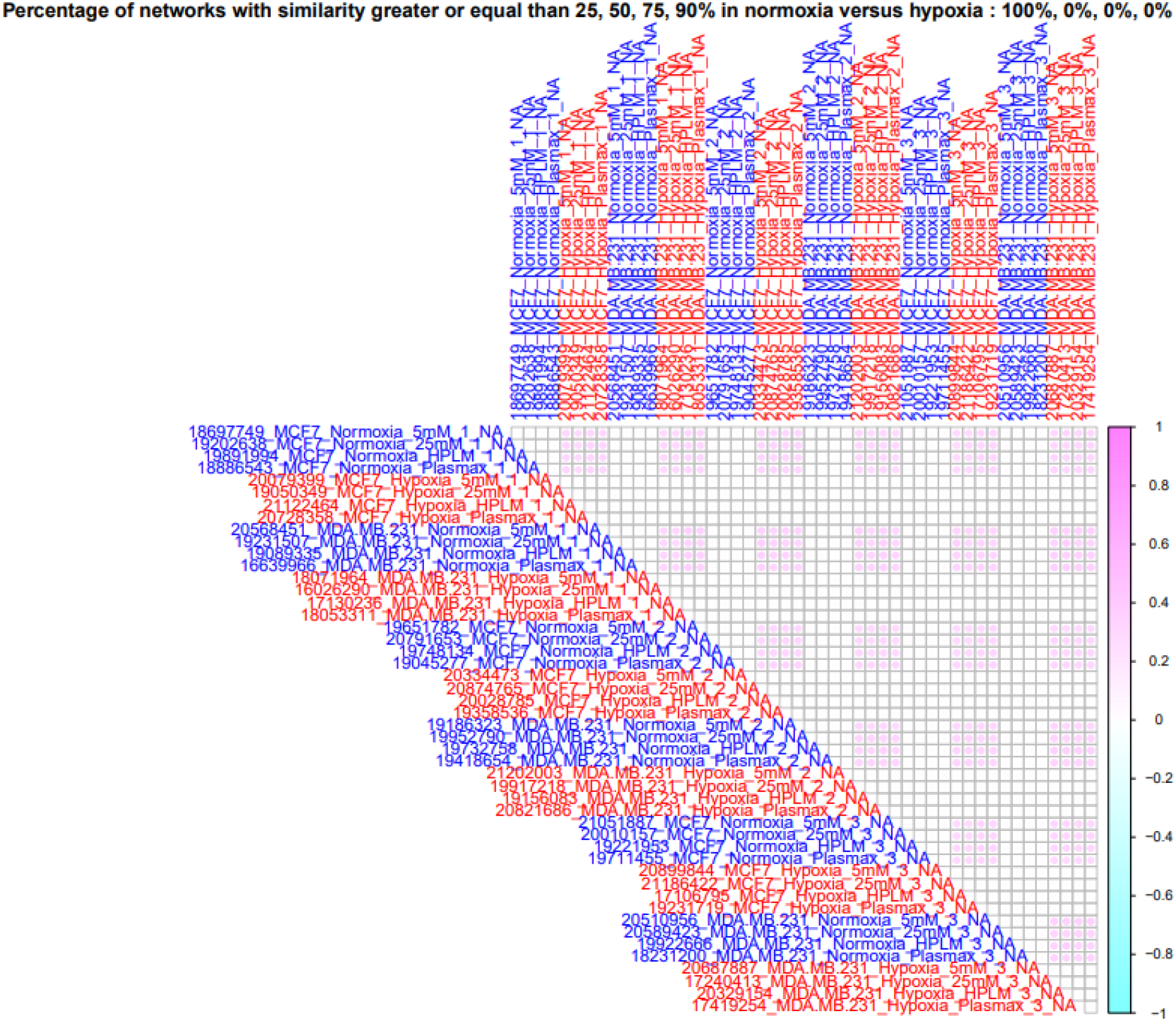
Comparison between normoxia and hypoxia only. As expected, the network pairs compared are scoring low in similarity, with 0% of the total number of network comparisons scoring above 50% similarity.

## Footnotes

2 for example, for 100 optimized networks, we need 4950 comparisons

3 DepMap, Broad (2020): DepMap Public. figshare.data-set doi:10.6084/m9.figshare.9201770.v2.

